# Evaluation of consensus strategies for haplotype phasing

**DOI:** 10.1101/2020.07.13.175786

**Authors:** Ziad Al Bkhetan, Gursharan Chana, Kotagiri Ramamohanarao, Karin Verspoor, Benjamin Goudey

## Abstract

**Motivation:** Haplotype phasing is a critical step for many genetic applications but incorrect estimates of phase can negatively impact downstream analyses. One proposed strategy to improve phasing accuracy is to combine multiple independent phasing estimates to overcome the limitations of any individual estimate. As such a strategy is yet to be thoroughly explored, this study provides a comprehensive evaluation of consensus strategies for haplotype phasing, exploring their performance, along with their constituent tools, across a range of real and simulated datasets with different data characteristics and on the downstream task of genotype imputation.

**Results:** Based on the outputs of existing phasing tools, we explore two different strategies to construct haplotype consensus estimators: voting across outputs from multiple phasing tools and multiple outputs of a single non-deterministic tool. We find the consensus approach from multiple tools reduces switch error by an average of 10% compared to any constituent tool when applied to European populations and has the highest accuracy regardless of population ethnicity, sample size, SNP-density or SNP frequency. Furthermore, a consensus provides a small improvement indirectly the downstream task of genotype imputation regardless of which genotype imputation tools were used. Our results provide guidance on how to produce the most accurate phasing estimates and the tradeoffs that a consensus approach may have.

**Availability:** Our implementation of consensus haplotype phasing, consHap, is available freely at https://github.com/ziadbkh/consHap.

## Introduction

Computational haplotype phasing, whereby phase information is statistically estimated from genotype data, remains an essential task in many types of genetic studies such as genome-wide association studies (GWAS) [1], expression quantitative trait loci (eQTL) [2] and genotype imputation [3, 4, 5]. While short-read sequencing technologies can provide phase information [6], studies based on SNP array remain more widely-used, hence there is still a strong demand for statistical approaches for estimating phase. An illustration of this demand is seen in the popular genotype imputation services provided by Sanger [7] and Michigan [4], where the latter has imputed over 58.1 million genomes before July 2020. In the imputations provided by these servers, users select a tool to use for phasing, which will in turn have an impact on the accuracy of all downstream analyses. Given its strong impact as a pre-processing step, it is critical to understand and improve the accuracy of approaches to phasing.

While state-of-the-art phasing tools have an error rate of only a few percent on large, high-density GWAS, error rates are significantly higher in cohorts with low sample size or low-SNP density [8, 9]. This can affect larger studies constructed from multiple smaller cohorts, such as the Haplotype Reference Consortium (HRC) dataset [7], which consists of 38,821 individuals from 20 different cohorts, with mean sample size is 1,630 individuals. Collating such data often requires genotype imputation for study harmonisation which is influenced by phasing accuracy within the small datasets.

One approach that has been proposed to improve phasing accuracy is to combine different estimations of haplotypes into a single consensus estimator through majority voting, a form of ensemble learning [10]. The different haplotypes can be obtained from either several tools [11, 12], which we denote a *multi-tool* consensus, or several iterations of a single but non-deterministic tool [13, 14, 9], which we denote a *multi-iteration* consensus. While these consensus or ensemble approaches have been previously mentioned, there has been little study on which phasing estimates to combine, how to combine them or the impact such techniques have on overall accuracy. Furthermore, preliminary analysis [11] and studies of ensemble methods in the machine learning community [15] highlight that combining multiple models does not necessarily outperform individual tools. Given a consensus estimator has an increased computational cost, requiring a dataset to be phased multiple times, a more rigorous study of this approach is warranted to understand its benefits and trade-offs.

This study provides an evaluation of consensus haplotype approaches. Several combinations of multiple tools or multiple iterations of the same tool are used to construct and evaluate the accuracy of this strategy and the results are compared to the constituent tools. Using data from HapMap III and HRC, we conducted a comprehensive evaluation of the proposed consensus estimators against state-of-the-art population-based phasing tools and focusing on the ones employed by Michigan and Sanger imputation servers (SHAPEIT2 [16], EAGLE2 [8], SHAPEIT3 [9], and HAPI-UR [14]). We evaluate how the proposed methods are affected by factors known to influence phasing accuracy including sample size, SNP density, minor allele frequency (MAF) and population ethnicity. We explore the impact of pre-phasing on genotype imputation across several imputation tools (pbwt [5] used in Michigan imputation server, Minimac3 [7] used in Sangar imputation server, and Beagle5 [3]). Finally, we analyse the trade-off between runtime and accuracy for the proposed methods and discuss several factors that are likely to mitigate the additional computational cost.

Our results show that consensus haplotype approaches are consistently the most accurate approach for phasing, with computational costs likely to be acceptable in most common scenarios. We also find that consensus haplotype phasing leads to the most accurate imputation compared to most combinations of phasing and genotype imputation tools available on Sanger and Michigan imputation servers, albeit with only modest increases over individual tools. SHAPEIT2 and EAGLE2 provide the most accurate phasing from any individual phasing tool, with EAGLE2 provided better genotype imputation in our evaluation. Our study results provide guidance on the strengths and limitations of consensus strategies for haplotype phasing that can help other researchers to conduct accurate genetic investigations and highlight several directions for future research.

## Methods

### Study workflow

Figure 1 illustrates the applied analysis workflow: **1)** Datasets preparation. **2)** Real haplotypes preparation. **3)** Phasing is applied to the Hap+HRC dataset using all individual phasing tools, then results are combined into a consensus estimators in 4). **5)** Estimated HAPMAP III Trios are evaluated against real haplotypes using switch error. **6)** The SNPs of the reference panel are imputed to the phased Hap+HRC dataset using three genotype imputation tools. **7)** Imputed genotypes are evaluated against the experimental genotypes filtered in step (1). **8)** Correlation assessment for phasing and genotype imputation accuracy at each individual scale (same individuals in step (5)).

**Figure 1:**
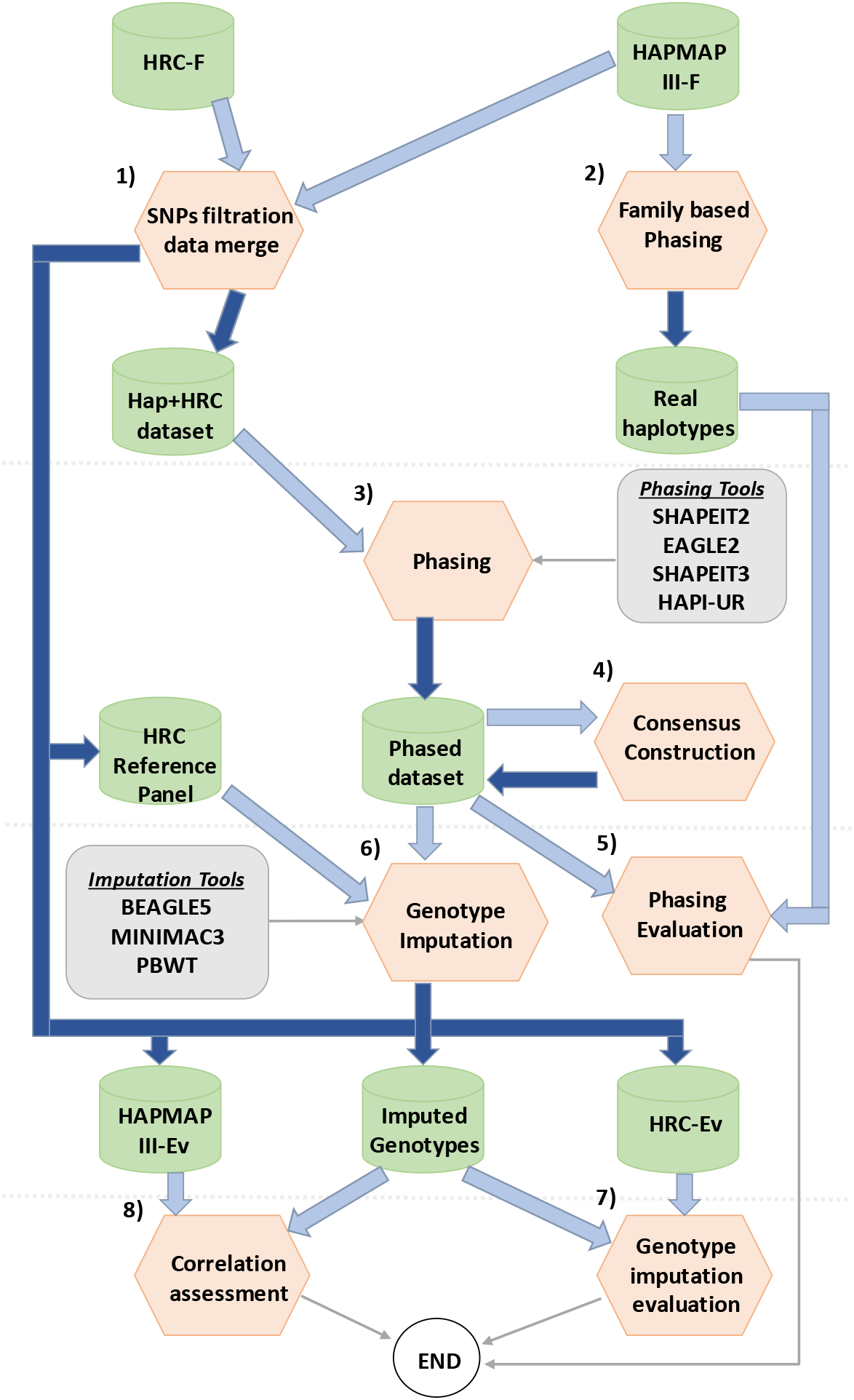
Workflow applied in this study. Dark blue arrows represent saving data into files, while light blue arrows represent reading the saved data. Thin arrow used to show additional information or the end of the workflow. Cylinders are used to represent data storage. Hexagons are used to represent applied procedures.

### Datasets and preparation

We analyzed data from HapMap phase III [17] and the Haplotype Reference Consortium (HRC) [7] with quality control for these datasets described in their respective publications. The majority of the analysis focuses on a combination of European samples from HapMap III and HRC datasets, which we denote the *Hap+HRC* dataset. We make use of 8,259 individuals of European descent, combining 69 Utah residents with Northern and Western European ancestry (CEU) obtained from HapMap phase III and 8,189 British samples from HRC (specifically the UK10UK and IBD cohorts).

HRC and HapMap samples were combined, keeping SNPs with MAF > 0.05 and are common to the two datasets. As both studies have very high density, we use LD-based pruning via PLINK [18] (–*indep-pairwise* with *r*^2^ = 0.95), resulting in a SNP density similar to the UK Biobank dataset as per [8]. Hap+HRC individuals with the original SNP density were saved in *HAPMAP III-Ev* and *HRC-Ev* for genotype imputation evaluation. All analysis for the Hap+HRC dataset focus on 5 chromosomes, totalling 151,278 SNPs (50,521 on chr2, 38,930 on chr6, 30,580 on chr16, 21,928 on chr6 and 9,319 on chr21). Phasing accuracy of the Hap+HRC dataset is benchmarked using 52 trios in the CEU dataset (denoted *Real haplotypes*), where phase is resolved using parental genotypes. As the HRC dataset consists of statistically estimated haplotypes, it could not be used for evaluation. HRC samples excluded from the Hap+HRC dataset are used as a reference panel for genotype imputation.

We additionally use all 838 samples (excluding any parental samples) from the HapMap phase III to emulate phasing a multi-ethnic cohort, using the same SNPs of chromosome 21 of the Hap+HRC dataset. Evaluation is conducted using trios from five populations: African ancestry in Southwest USA (ASW), Utah residents with European ancestry (CEU), Mexican ancestry in Los Angeles (MEX), Maasai ancestry in Kenya (MKK), and Yoruba in Ibadan, Nigeria (YRI). The number of trios in each population is reported in Table 2.

### Haplotype phasing and genotype imputation tools

Haplotype phasing was carried out using four tools: SHAPEIT2 (v2.r904), EAGLE2 (v2.4.1), HAPI-UR (v1.01), and SHAPEIT3 (v3.r881), with the first three tools motivated by their availability as part of the widely-used Sanger and Michigan genotype imputation servers. Phasing tools were applied using their default parameters unless otherwise was noted. SHAPEIT3 was applied with fast flag enabled as mentioned in its documentation. When phasing small datasets via SHAPEIT3, the cluster size parameter was set to (2× sample size - 1), a mandatory setting when the sample size is below 4,000.

We utilised the following genotype imputation tools applied with default parameters: Minmac3 (2.0.1), used in the Michigan imputation servers, pbwt (3.0), used in the Sanger imputation servers, and Beagle v5.

### Consensus estimator

Rather than making use a single model, a consensus approach is a form of ensemble learning, whereby the predictions from multiple independent models are combined to obtain better predictive performance than any of the constituent models [10]. In the context of haplotype phasing, existing phasing algorithms are usually based on Hidden Markov Models, with the different tools each making different assumptions and implementing different heuristics to improve runtime [19, 13]. By combining the predictions of these models, it is hoped that a majoity vote can overcome the limitations and biases of each individual model, resulting in a higher accuracy on average.

IN this work, we focus on consensus approach, demonted *consHap*, based on a majority voting approach described in Algorithm 1 and previously proposed in [11, 20]. The method takes *M* phasing estimates where the output from the *j*-th tool, denoted *h_j_*, is a binary array of length *P*, representing the sequence of alleles at every heterozygous SNP on a single homologous chromosome. hj [i] denotes whether the i-th heterozygous SNP carries a major or minor allele (0 or 1 respectively). Only one homologous chromosome needs to be represented, with the remaining copy being the logical complement. For each SNP, our method takes a majority vote across all input phasing estimates. Any method that is not concordant with the majority vote has phasing estimates for remaining SNPs flipped (i.e we take the logical complement) in order to align the phase of the remaining estimates with those of the majority.

#### Algorithm 1: Consensus haplotype estimator algorithm.

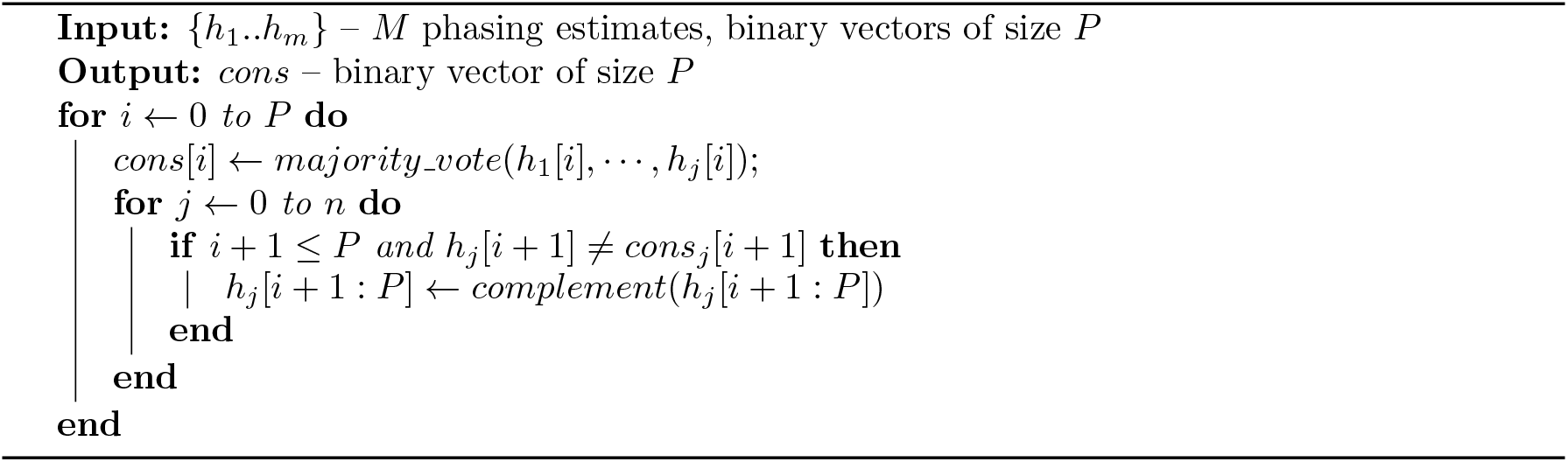

We consider two different forms of consensus estimators, either multi-tool approaches that combine the output of different phasing tools, or multi-iteration approaches that combine the output of multiple iterations of one non-deterministic tool. We also explore the impact of combining these approaches i.e. multiple iterations from multiple tools. Most phasing tools considered in this work are non-deterministic, in the sense that they produce different outputs when running with the same data. Only EAGLE2 produces deterministic results but can be forced to emulate non-deterministic output by permuting the order of individuals in the input [11].

We use a consistent naming convention to describe a particular implementation of consHap throughout this work. Individual tools are abbreviated (*S*_2_: SHAPEIT2, *S*_3_: SHAPEIT3, *E*_2_: EAGLE2, *H_r_*: HAPI-UR) and concatenated together if multiple tools are used. If multiple outputs from a single tool are used, we denote the number of iterations in parentheses. Thus consHap-*S*_2_*E*_2_*S*_3_ is based on the output of SHAPEIT2, EAGLE2, and SHAPEIT3 while consHap-*S*_2_ *(3)* is based on three different iterations of SHAPEIT2.

### Evaluation criteria

The evaluation of phasing accuracy is based primarily on the proportion of switch errors (SE), which occur when the phase allocated to a heterozygous variant is incorrectly switched relative to the prior heterozygous [13]. While switch error rate is primarily reported as an average across all samples in a dataset, we also consider the variation of switch error at an individual level. This also enables us to evaluate there is a statistically significant difference between a pair of phasing tools via a one-sided binomial test, with the number of trials as the number of individuals and the probability of an improvement as 0.5 given the null hypothesis that two tools have the same accuracy on average.

Evaluation of phasing performance with varying SNP density was conducted on five subsets of the Hap+HRC datasets, restricted to chromosomes 16 and 21, and thinned to a SNP density ranging from 200SNPs/1Mb to 1000 SNPs/Mb. This pruning (via PLINK’*s* –*bp-space* command) removed a random SNP from any possible pair if the distance was less than the desired resolution. Similarly, five datasets with different population sizes were generated by a random selection of samples from the Hap+HRC dataset, again restricted to chromosomes 16 and 21. The error rate of phasing as MAF varies was calculated across all chromosomes of the Hap+HRC dataset.

Using the Hap+HRC dataset that was thinned for the phasing evaluation, we imputed all pruned SNPs using the reference panel into the Hap+HRC dataset, a masking experiment similar to [21]. Concordance of imputed and actual genotype calls was measured using Pearson correlation between the allele dosage and the sum of the posterior allele probabilities 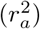 for each SNP [3, 4].

## Results

### Consensus performance across European cohort

We first evaluated consensus haplotype approaches, which we refer to as *consHap*, across five chromosomes of the Hap+HRC dataset and compared the proportion of switch errors (SE) with that achieved by several state-of-the-art phasing tools. We focus on two variants: a multi-tool consensus of SHAPEIT2, EAGLE2 and SHAPEIT3 (consHap-*S*_2_*E*_2_*S*_3_) and a multi-iteration consensus of three outputs of SHAPEIT2 (consHap-*S*_2_(3)).

Table 1 shows that the consensus estimators, consHap-*S*_2_*E*_2_*S*_3_ and consHap-*S*_2_(3), improved on the most accurate individual tool across the evaluated chromosomes of the Hap+HRC dataset by an average of 9.8% (consHap-*S*_2_*E*_2_*S*_3_) and 5.6%(consHap-*S*_2_(3)). The multi-tool consHap-*S*_2_*E*_2_*S*_3_ consistently improves upon the multi-iteration consHap-*S*_2_(3) by between 2.3 and 8.8% depending on the chromosome. SHAPEIT2 is the most accurate individual tool in all cases, showing consistent improvements beyond EAGLE2. The average length of correctly phased haplotype blocks obtained by cons-S2E2S3 and cons-S2(3) is 421, and 403 SNPs respectively, representing 45 and 27 SNPs longer blocks compared to the best individual tool as shown in supplementary Table 3.

**Table 1:** Switch error % and total number of switches for evaluated phasing tools on 52 individuals for five chromosomes of the Hap+HRC dataset. Bolded values represent the lowest error while underlined values have the lowest switch error for any individual tool (i.e excluding consensus). Improvement (bottom row) highlights the percentage of switch error improvement between consHap-*S*_2_*E*_2_*S*_3_ and the best individual tool.

| Tool | Chr2 | Chr6 | Chr11 | Chr16 | Chr21 | Switches |
| --- | --- | --- | --- | --- | --- | --- |
| **consHap-$S_2E_2S_3$** | **0.86** | **0.83** | **0.92** | **1.10** | **1.23** | **18,763** |
| consHap-$S_2(3)$ | 0.88 | 0.85 | 0.95 | 1.14 | 1.32 | 19,392 |
| **SHAPEIT2** | <u>0.95</u> | <u>0.92</u> | <u>1.00</u> | <u>1.20</u> | <u>1.40</u> | <u>20,742</u> |
| **EAGLE2** | 0.99 | 0.94 | 1.04 | 1.24 | 1.41 | 21,352 |
| **SHAPEIT3** | 1.1 | 1.06 | 1.16 | 1.38 | 1.51 | 23,850 |
| **HAPI-UR** | 1.51 | 1.42 | 1.61 | 1.89 | 2.12 | 32,604 |
| **Improvement** | 9.76% | 10.38% | 8.15% | 8.52% | 11.72% | 1,979 |

**Table 2:**
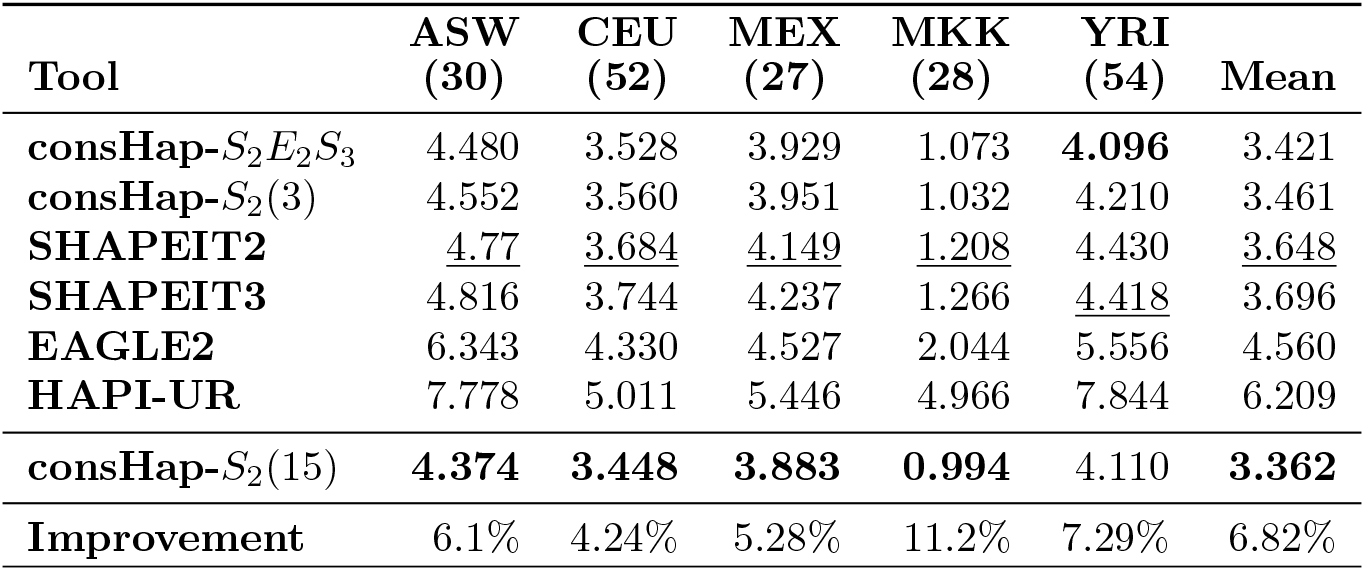
Switch error % when phasing different populations from the HapMap III project. The title of each column shows abbreviated population names and number of trios (in parentheses), with the full populations names in Section. Populations information are in supplementary methods. The approaches are sorted based on the average switch errors. Bolded results are the lowest switch error. Italic results are the best across all tests. Underlined results are the best individual tool. The improvement row demonstrates the percentage improvement between consHap-*S*_2_*E*_2_*S*_3_ and the best single tool.

Rather than looking at average switch error, we can also consider how often a given phasing tool results are the most accurate phasing of an individual. Supplementary Table 2 shows that consHap-*S*_2_*E*_2_*S*_3_ and consHap-*S*_2_(3) were more accurate than all individual tools for 76% and 56% of individuals, representing a significant improvement in performance (one-sided binomial test *p* < 5 × 10^−5^ for both consensus methods). The scenario where individual tools perform better than the consensus arises when the majority of phasing outputs in the consensus are in error but out-vote the more accurate minority.

We also explored a range of other consensus configurations (Supplementary Table 4 and 5). The highest accuracy was obtained by consHap-*S*_2_(8)*E*_2_(7), a consensus of 8 iterations of SHAPEIT2 combined with 7 iterations of EAGLE2, showing a mean SE of 1.057% compared to 1.168% by consHap-*S*_2_*E*_2_*S*_3_, but with a cost of a significant increase in runtime. Supplementary Table 4 shows that consensus estimators constructed from any three tools obtained higher accuracy than any of the tools individually. However, Supplementary Figure 1 shows this was not the case with a multi-iteration consensus, where multiple iterations of SHAPEIT3 or HAPI-UR failed to outperform a single iteration of SHAPEIT2 or EAGLE2.

### Performance across factors that influence accuracy

We evaluated consHap-*S*_2_*E*_2_*S*_3_ and consHap-*S*_2_ (3) over several subsets of the Hap+HRC dataset to assess their robustness against factors known to influence phasing accuracy including SNP density, population size, and minor allele frequency (MAF) [22, 13].

As shown in Figure 2, consHap, in both multi-tool and multi-iteration configurations, outperforms all individual tools regardless of SNP density from subsampled versions of the Hap+HRC dataset. Overall, the average SE increases by an average of 1.94% to 4.2% across all approaches as the SNP density is reduced. When changing the resolution of SNPs from 200SNPs/Mb to 1000SNPs/Mb, as shown in Figure 2 (a), we see consHap-*S*_2_*E*_2_*S*_3_ and consHap-*S*_2_(3) improved the accuracy by an average of 8.54% and 6.65% respectively compared to the best single tool. Interestingly, the best individual tool changes from SHAPEIT2 to EAGLE2 as the density of SNPs gets higher than 500SNPs/Mb. We again observe that the multi-tool consensus, consHap-*S*_2_*E*_2_*S*_3_, outperforms the multi-iteration consensus, consHap-*S*_2_(3) showing a mean improvement of 2%.

**Figure 2:**
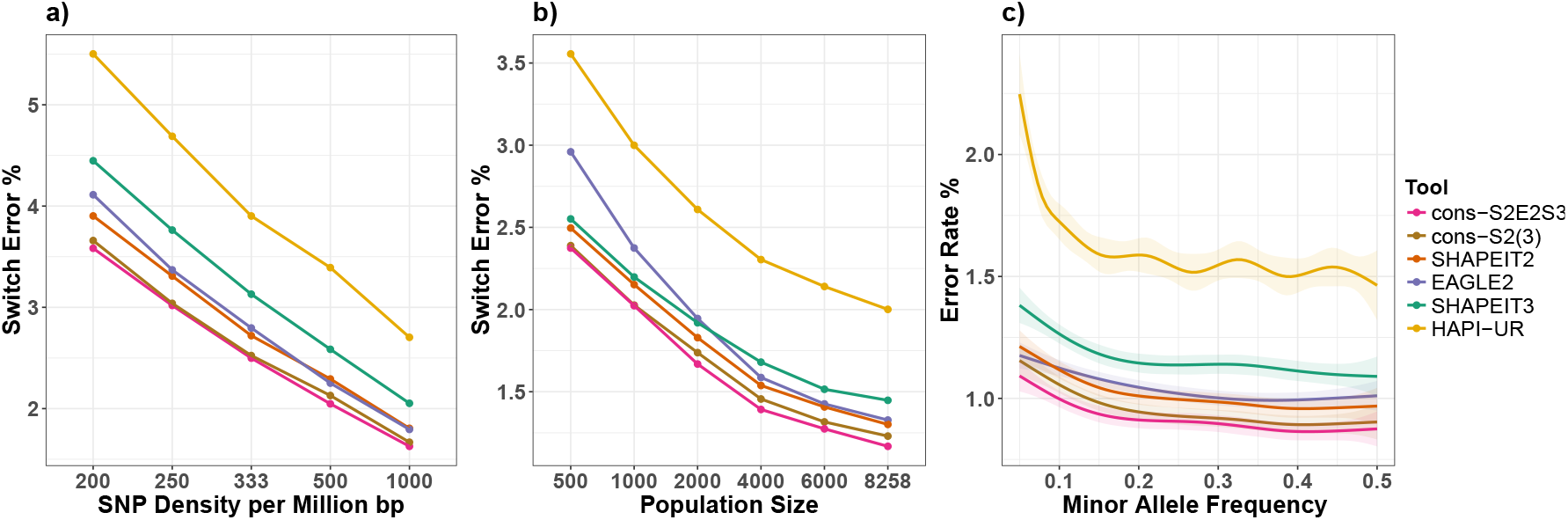
Robustness of the consensus approach to a) SNP density, b) population size and c) minor allele frequency (MAF), as measured by switch error and Error rate. c) is generated from the Hap+HRC dataset and smoothed using a generalized additive model (GAM), while a) and b) are generated by manipulating the data of chromosomes 16 and 21 from the Hap+HRC dataset to obtain the desired characteristics as explained in supplementary materials.

Similar results are seen when varying sample size. In Figure 2 (b) we see consHap-*S*_2_*E*_2_*S*_3_ and consHap-*S*_2_(3) improve accuracy by 7.7% and 4.4% when changing the population size from 500 to 8,258 individuals, the percentage improvement increases as sample size increases from 4.8% to 11.7% and 3.5% to 5.4% for consHap-*S*_2_*E*_2_*S*_3_ and consHap-*S*_2_(3), respectively.

Furthermore, consHap obtained the minimal error rate regardless of the minor allele frequency (MAF) as illustrated in Figure 2 (c). There was 9.7% (averaged for all MAF ranges) error reduction obtained by consHap-*S*_2_*E*_2_*S*_3_ compared to the best individual tool. While the accuracy of all phasing tools decreases as we examine increasingly rare SNPs, we observe that while SHAPEIT2 is the most accurate individual tool for variants with MAF > 0.1, its accuracy drops below that of EAGLE2 once when MAF < 0.1.

### Accuracy gains of increasing consensus iterations

The results in previous sections consider consHap-*S*_2_(3), a specific instance of consHap-*S*_2_(*n*), a consensus estimator based on *n* iterations of SHAPEIT2. We can evaluate the impact on performance as the parameter for iterations, *n,* is increased. Moreover, as the consensus relies on the non-deterministic output of SHAPEIT2, we can also examine the distribution of accuracy from different runs of this tool. This is achieved by running SHAPEIT2 up to 50 times and, for each *n* of interest, selecting *n* random iterations as inputs into the multi-iteration consensus. This selection is repeated 10 times, to derive the distribution of error.

**Figure 3:**
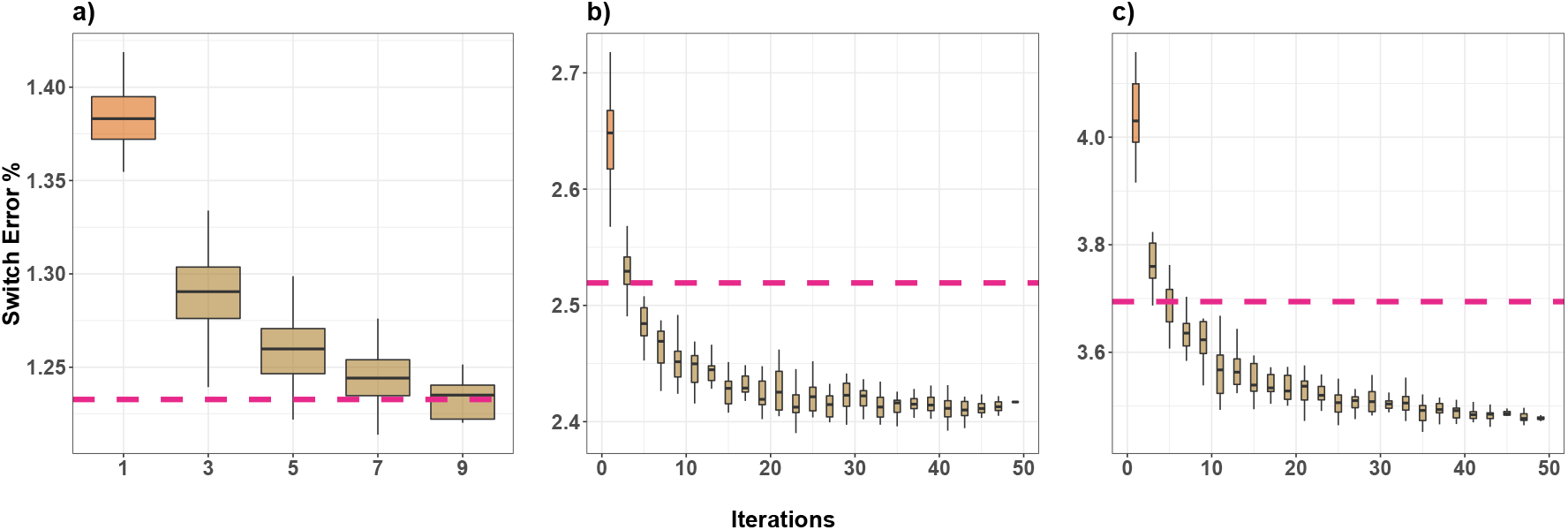
Performance of multi-iteration consensus for varying numbers of iterations over chromosome 21 of the Hap+HRC dataset a) without modification, b) reduced to 500 individuals and c) with SNP density reduced to 1 SNP per 5kb. The number of iterations is shown on the x-axis. The dashed pink line shows switch error for consHap-*S*_2_*E*_2_*S*_3_.

Initially, we examine the performance of consHap-*S*_2_ (*n*) as *n* is increased, focusing on only chromosome 21 of the Hap+HRC dataset. Figure 3 (a) highlights a 7.2% improvement in accuracy when shifting from 1 iteration to 3 iterations (mean SE from 1.39% to 1.29%, respectively) with 9 iterations required to achieve the same mean SE as the multi-tool consHap-*S*_2_*E*_2_*S*_3_ (SE = 1.23%). Additional iterations continue to reduce SE with diminishing returns, with details of the accumulative improvement is reported in Supplementary Table 6. Similar observations are noted when constructing a multi-iteration consensus from other tools (Supplementary Figure 1).

As datasets with low sample size or SNP density can be phased faster, we hypothesised that a multi-iteration consensus combining a large number of phasing outputs may outperform a multitool consensus. Evaluating up to 49 iterations on a data set with reduced sample size (n=500), we found that consHap-S2(3) obtained similar results to consHap-S2E2S3 (SE = 2.53%) as illustrated in Figure 3 (b). Similar results are observed when SNP density is reduced but sample size is maintained, as shown in Figure 3 (c). Here, consensus from five iterations of SHAPEIT2 or more improve beyond consHap-*S*_2_*E*_2_*S*_3_ (SE = 3.7%). While these results were produced for chromosome 21, Supplementary Figure 2 shows similar findings on chromosome 16. Supplementary Figure 3 shows these results hold as population size and SNP density are varied further.

To understand why these multi-iteration consensuses improve beyond consHap-S2E2S3, we in-vestigated the individual tools used in the consensus construction. Consistent with results in Figure 2, Supplementary Figure 4 shows EAGLE2 has a relatively high SE over datasets with small sample size (SE= 3.2% compared to 2.64% and 2.69% for SHAPEIT2 and SHAPEIT3, respectively) that may impact the performance of consHap-*S*_2_*E*_2_*S*_3_ in Figure 3 (b). When SNP density is reduced, SHAPEIT3 obtained SE = 4.6% compared to 4.17% and 4.04% for EAGLE2 and SHAPEIT2 respectively, which may impact consHap-S2 E2 S3 in Figure 3 (c). These analyses indicates that SHAPEIT2 has the most robust performance across low sample and low SNP density datasets. Additionally, we found that constructing a consensus using several iterations of one non-deterministic tool is not guaranteed to outperform all individual tools, with Supplementary Figure 5 showing multiple iterations of HAPI-UR failing to outperform a single iteration of SHAPEIT2 or EAGLE2, given the large gap in accuracy.

### Phasing performance on a multi-ethnic cohort

We evaluate the performance of phasing on a multi-ethnic cohort formed by combining all samples from the HapMap. As shown in Table 2, we find consHap-*S*_2_*E*_2_*S*_3_ outperforms all individual tools, with improvements ranging from 4.2% (CEU) to 11.2% (MKK). consHap-*S*_2_*E*_2_*S*_3_ also outperforms consHap-*S*_2_(3) across all populations, though the improvement is only 1.15% on average.

As the smaller sample size used for this experiment means error rates across all phasing tools are typically high (3-4%) for all ethnicities, we see a lower error rate in the Maasai in Kinyawa, Kenya (MKK) cohort, likely because this is the largest population in this multi-ethnic dataset. We also observe a substantially higher error rate for the CEU trios compared to Table 1, due to both the smaller number of samples used for phasing and the non-European nature of this experiment.

Given the results from Figure 3 showing that higher iterations of a single tool had stronger results than a multi-tool consensus, we repeated the analysis with 15 iterations of SHAPEIT2 (consHap-*S*_2_(15)). The resulting error rates were lower than all other phasing estimators, with the exception of the YRI population where consHap-*S*_2_*E*_2_*S*_3_ was the most accurate approach.

### Impact of different phasing strategies on genotype imputation

Evaluations of phasing reported in this work consistently show the improvements obtained by consHap. However, we can also assess the impact of consHap on downstream tasks. We consider the use of consHap as well as all phasing tools for pre-phasing for genotype imputation. As SNPs were filtered out of Hap+HRC, we imputed them again and compared them to the real SNPs obtained experimentally with the original dataset. Genotype imputation tools (Beagle5, Minimac3, and pbwt) are applied to the Hap+HRC dataset after phasing by all methods described so far, as well as considering the data unphased.

The results in Table 3 for SNPs within three ranges of MAF show imputation using Beagle5 consistently obtained the highest accuracy, followed by Minimac3 and pbwt. With respect to phasing tools, consHap, regardless of configuration, obtained higher accuracy than any individual tool, with the best results obtained by consHap-*S*_2_(8)*E*_2_(7). Compared to the results obtained by the best individual tool for pre-phasing, consHap-*S*_2_(8)*E*_2_(7) and consHap-*S*_2_*E*_2_*S*_3_, correctly imputed a further 2,800,715 and 1,266,483 genotypes respectively, which equates to 350 correctly imputed SNPs for each individual, within 5 chromosomes. The highest relative improvement was for SNPs with MAF in the range from 0.5% to 5%. With regard to pre-phasing performed via a single tool, EAGLE2 had the highest accuracy in all but two cases where SHAPEIT2 was better. Detailed results showing further imputation metrics are shown in Supplementary Tables 7-9.

Given the results in Table 3 appear to indicate that improved phasing accuracy may have little improvement in genotype imputation, we sought to quantify this relationship. The correlation between phasing and genotype imputation accuracy was evaluated for the 52 individuals of HapMap III, as these individuals have both known real haplotypes and high-density genotype data. Results in Supplementary Figures 6 and 7 show a positive correlation between phasing and imputation accuracy with an mean Pearson correlation across all individuals of r=0.35 (range: 0.26 to 0.48), a highly significant relationship (mean *p* = 6 × 10^−6^, range: 6 × 10^−16^ to 3 × 10^−5^). Overall, Beagle5 was more influenced by phasing accuracy compared to other imputation tools mean r = 0.4, mean p = 4 × 10^−9^), then Minimac3 (mean *r* = 0.35, mean p = 9 × 10^−7^) and finally pbwt (mean *r* = 0.3, mean p = 7.1 × 10^−6^). Detailed results are reported in Supplementary Table 10.

**Table 3:** Genotype imputation evaluation. r^2^ summarised for 54,230,334,071 SNPs across three MAF ranges: 0.01% to 0.5%, 0.5% to 5%, and 5% to 50%. Improvement column represents the accumulative additional correctly imputed SNPs compared to the pre-phasing done by the phasing tool in the next row. For example, the additional correctly imputed SNPs by consHap-*S*_2_(8)*E*_2_(7) in the first row is 1,447,183 SNPs compared to consHap-*S*_2_*E*_2_*S*_3_ and (1,447,183 + 560,920) SNPs compared to consHap-*S*_2_(3). Bold results indicate the highest accuracy for each imputation tool, underlined results indicate the most accurate individual phasing tools.

|  | Phased by | [0.01%,0.5% ] | (0.5%, 5%] | ≥ 5% | Improvement |
| --- | --- | --- | --- | --- | --- |
| Imputed by Beagle5 | consHap-$S_2(8)E_2(7)$ | **0.314** | **0.768** | **0.972** | 1,447,183 |
| | consHap-$S_2E_2S_3$ | 0.309 | 0.766 | 0.971 | 560,920 |
| | consHap-$S_2(3)$ | 0.308 | 0.765 | 0.971 | 723,566 |
|  | EAGLE2 | <u>0.307</u> | <u>0.763</u> | <u>0.971</u> | 254,420 |
|  | SHAPEIT2 | 0.306 | <u>0.763</u> | <u>0.971</u> | 2,817,004 |
|  | SHAPEIT3 | 0.299 | 0.758 | 0.97 | 3,740,168 |
|  | HAPI-UR | 0.296 | 0.751 | 0.969 | 1,032,159,933 |
|  | unphased | 0.066 | 0.327 | 0.745 | - |
| Imputed by Minimac3 | consHap-$S_2(8)E_2(7)$ | **0.315** | **0.764** | **0.971** | 1,473,243 |
| | consHap-$S_2E_2S_3$ | 0.31 | 0.762 | 0.97 | 545,408 |
| | consHap-$S_2(3)$ | 0.309 | 0.761 | 0.97 | 660,184 |
|  | EAGLE2 | <u>0.308</u> | <u>0.759</u> | <u>0.97</u> | 285,388 |
|  | SHAPEIT2 | 0.307 | <u>0.759</u> | <u>0.97</u> | 2,887,572 |
|  | SHAPEIT3 | 0.299 | 0.754 | 0.969 | 355,6971 |
|  | HAPI-UR | 0.297 | 0.746 | 0.969 | 980,415,942 |
|  | unphased | 0.065 | 0.298 | 0.759 | - |
| Imputed by pbwt | consHap-$S_2(8)E_2(7)$ | **0.279** | **0.739** | **0.969** | 1,682,270 |
| | consHap-$S_2E_2S_3$ | 0.274 | 0.736 | 0.968 | 723,411 |
| | consHap-$S_2(3)$ | 0.272 | 0.735 | 0.968 | 585,959 |
|  | EAGLE2 | <u>0.271</u> | <u>0.734</u> | <u>0.968</u> | 541,770 |
|  | SHAPEIT2 | 0.27 | 0.733 | <u>0.968</u> | 3,281,317 |
|  | SHAPEIT3 | 0.261 | 0.727 | 0.967 | 4,157,745 |
|  | HAPI-UR | 0.259 | 0.719 | 0.966 | 1,140,380,286 |
|  | unphased | 0.047 | 0.249 | 0.725 | - |

### Runtime and computational cost

The improved accuracy of the consensus comes at a cost of increased runtime. Figure 4 shows the runtime of the individual tools used in this analysis when evaluated across chromosome 21 of the Hap+HRC dataset, as the SNP density and sample size are varied. When phasing tools are applied in parallel, the total time required by consHAP can be calculated as *max*(*t*_1_,*t*_2_, …, *t_n_*) + *t*_0_ where *t_n_* is phasing time of the tool *n* and *t*_0_ is the time required to aggregate the results together. If the tools are applied sequentially, the total time can be calculated as 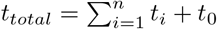. For chromosome 21 of the Hap+HRC dataset, it required 32.2 and 42.6 hours to construct consHap-S2E2S3 and consHap-S2 (3) considering the worst-case scenario where phasing is applied sequentially and using one thread. This runtime reduces dramatically when phasing tools can be run in parallel to 14.2 hours for both consHap-*S*_2_*E*_2_*S*_3_ and consHap-*S*_2_(3), with SHAPEIT2 as the bottleneck. The time to conduct voting across the tools was 1.6% of the time to run the tools in parallel (6 minutes). Individually, HAPI-UR was the fastest phasing tool, with a required execution time around 11 times faster than the closest competitor EAGLE2, while SHAPEIT2 is the slowest. Most tools scale linearly with the number of samples, with the exception of SHAPEIT2, while the high speed of phasing for HAPIUR compared to other approaches is demonstrated across all SNP densities and sample sizes.

**Figure 4:**
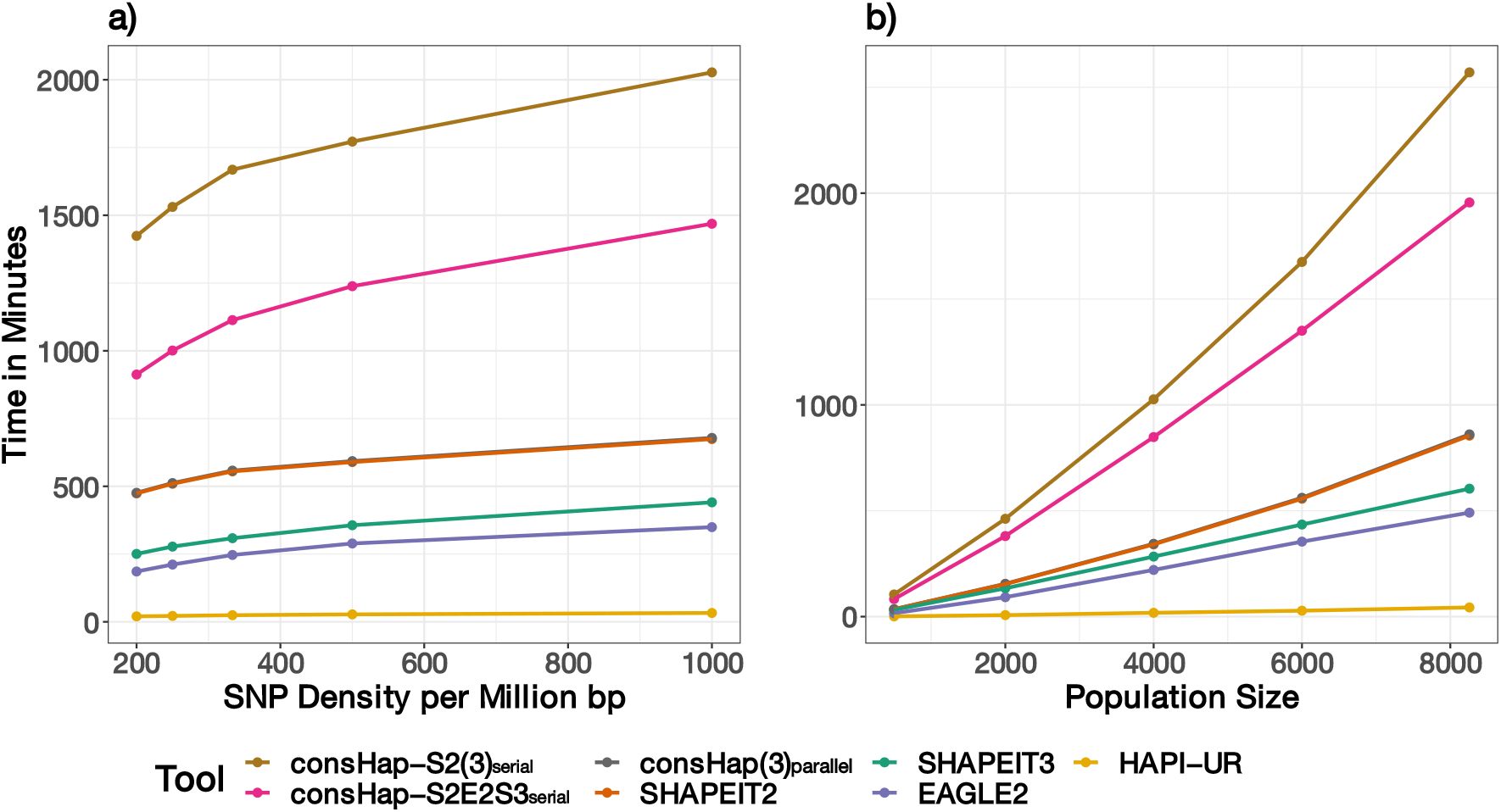
The execution time of phasing tools with respect to population size in (a) and SNP density in (b). Note we include timings for consHap when tools are run in serial or parallel. Subsampling of SNPs and samples was performed as per Figure 3.

## Discussion

In this study, we evaluate the performance of consensus haplotype estimators that use majority voting to combine multiple phasing outputs from different phasing tools or multiple iterations of a single non-deterministic tool. This study provides the first comprehensive evaluation of consensus phasing approaches, demonstrating that the proposed consensus approach leads to stronger phasing accuracy compared to using a single tool and it is robust across many characteristics known to impact phasing performance. We also assess the impact of phasing accuracy on genotype imputation, one of the major application of phasing, across all phasing and genotype imputation tools available in the widely-used Sanger and Michigan imputation servers, with the resulting findings providing guidance for researchers using these services.

In our evaluations, a consensus from multiple phasing tools almost always has higher accuracy than a consensus from the multiple outputs from the same tool. The consensus estimators are a form of ensemble learning whereby multiple predictive models are combined to produce a single prediction. Ensemble models typically have the strongest improvement above their constituent predictors when each model provide accurate but independent predictions [10]. With phasing, we would intuitively imagine that phasing outputs from the same non-deterministic tool will be less independent than outputs from different tools. Only across datasets with low sample size or low SNP density did we observe improvements from combining multiple outputs of SHAPEIT2 compared to multi-tool consensus. Even here, we required at least 5 independent runs of SHAPEIT2 to see such improvements and improvements in SE remained less than 0.2%.

Our study also builds upon previous benchmarks of phasing tools to characterize the performance of the constituent tools. As with previous studies, we find no individual tool is the best in all scenarios. SHAPEIT2 and EAGLE2 provided the strongest results, while HAPI-UR performs consistently worse, as previously reported [11, 20]. Also consistent with prior results [8], SHAPEIT3 tended to provide somewhat less accurate results than SHAPEIT2 or EAGLE2 but had a reduced runtime and still had higher accuracy than HAPI-UR. One interesting observation was that EAGLE2 displayed a substantially sharper drop in accuracy when sample size was low (less than 2000 individuals). This may be due to EAGLE2 use of long-range phasing to generate an initial haplotype set when no reference panel is provided, but if only relatively few samples are present, it may be hard to find individuals who share haplotypes [13].

The improvements in phasing accuracy achieved through a consensus approach do so at the cost of extra computational overhead. However, there are three complementary aspects related to phasing runtime that mitigate the computational overhead incurred by our consensus approach. First, the strong focus on improving run-time for many phasing algorithms has led to computational complexity reducing from quadratic in the number of samples to linear [19]. Secondly, access to high-throughput computational resources, whether via local high-throughput compute servers or whether via cloud services, are being more accessible, enabling the parallel phasing of data. If the running of multiple phasing tools can be parallelized, the observed runtime will be approximately the runtime of the slowest individual tool. Finally, phasing typically only needs to be conducted once for a study as opposed to being an analysis that needs to be run multiple times, and hence the once-off additional costs, may not be substantial compared to its re-use as part of downstream analyses. While performance time reported in the results of this work represent the worst-case scenario where phasing was applied sequentially using one thread on an average PC, these three facets of computational performance indicate consensus phasing is likely to be a practical option for most datasets.

Our study makes an important contribution in demonstrating the impact of phasing error on genotype imputation. In line with previously reported results [4, 3], we found that genotype imputation tools obtain similar accuracy, with Beagle5 consistently reporting the highest accuracy regardless of variant allele frequency. For each imputation tool, we found that the improvement of phasing accuracy reported for the consensus estimators led to the most accurate genotype imputation with 2,800,715 and 1,266,483 additional correctly imputed SNPs for 8,189 individuals within 5 chromosomes when using consHap-*S*_2_(8)*E*_2_(7) and consHap-*S*_2_*E*_2_*S*_3_, respectively compared to the best individual tool. Errors in imputed genotypes of SNPs have been reported to increase false positives and negatives in downstream analysis [23]. Therefore, the improvements in accuracy obtained by the consensus estimator will contribute to more accurate genetic analyses.

A limitation of our phasing evaluation is that it was carried out on at most 191 individuals from HapMap III, with most analyses focusing on the subset of 52 CEU individuals. However, known haplotype data is not readily available in many large studies and many previous empirical evaluations also suffer from this constraint [13, 8, 9, 11]. We restricted our experiments to a set of phasing and imputation tools (using their default parameters) available in Michigan and Sanger imputation servers given their widespread usage and demonstrated utility. While this limitation means we do not explore some of the most recently published phasing tools, including SHAPEIT4 [6] and BEAGLE5 [3], the tools have tended to improve scalability substantially but have modest impacts on accuracy. As the computational complexity of phasing algorithms drops, we believe the case for consensus approaches will be strengthened, given increased runtime is the strongest drawback.

While the results in this work show the promise of consensus approaches to phasing, future investigations on how best construct a consensus estimator, such as weighting the output of each tool depending on the properties of the data being phased, may yield even stronger improvements in performance. It may also be useful to explore consensus constructed from more diverse tools, with novel approaches such as the random-forest based phasing provided by [24], potentially resulting in greater independence between inputs and hence likely producing a more accurate phasing result. And finally, it may also be possible to alter the optimization of existing HMM-based phasing tools so that they produce multiple local optima, rather than the single best solution, and take a vote over these without requiring multiple independent runs of the tool and hence producing a census result without the runtime cost.

## Acknowledgements

The Haplotype Reference Consortium (HRC) dataset is used in a form agreed by The University of Melbourne with Wellcome Trust Sanger Institute.

## References

[1] Brian L Browning and Sharon R Browning. Efficient multilocus association testing for whole genome association studies using localized haplotype clustering. Genetic Epidemiology: The Official Publication of the International Genetic Epidemiology Society, 31(5):365–375, 2007.

[2] Ryan Tewhey, Vikas Bansal, Ali Torkamani, Eric J Topol, and Nicholas J Schork. The importance of phase information for human genomics. Nature Reviews Genetics, 12(3):215, 2011.

[3] Brian L Browning, Ying Zhou, and Sharon R Browning. A one-penny imputed genome from next-generation reference panels. The American Journal of Human Genetics, 103(3):338–348, 2018.

[4] Sayantan Das, Lukas Forer, Sebastian Schönherr, Carlo Sidore, Adam E Locke, Alan Kwong, Scott I Vrieze, Emily Y Chew, Shawn Levy, Matt McGue, et al. Next-generation genotype imputation service and methods. Nature genetics, 48(10):1284, 2016.

[5] Richard Durbin. Efficient haplotype matching and storage using the positional burrows-wheeler transform (pbwt). Bioinformatics, 30(9):1266–1272, 2014.

[6] Olivier Delaneau, Jean-François Zagury, Matthew R Robinson, Jonathan L Marchini, and Emmanouil T Dermitzakis. Accurate, scalable and integrative haplotype estimation. Nature communications, 10(1):1–10, 2019.

[7] Shane McCarthy, Sayantan Das, Warren Kretzschmar, Olivier Delaneau, Andrew R Wood, Alexander Teumer, Hyun Min Kang, Christian Fuchsberger, Petr Danecek, Kevin Sharp, et al. A reference panel of 64,976 haplotypes for genotype imputation. Nature genetics, 48(10):1279, 2016.

[8] Po-Ru Loh, Petr Danecek, Pier Francesco Palamara, Christian Fuchsberger, Yakir A Reshef, Hilary K Finucane, Sebastian Schoenherr, Lukas Forer, Shane McCarthy, Goncalo R Abecasis, et al. Reference-based phasing using the haplotype reference consortium panel. Nature genetics, 48(11):1443, 2016.

[9] Jared O’Connell, Kevin Sharp, Nick Shrine, Louise Wain, Ian Hall, Martin Tobin, Jean-Francois Zagury, Olivier Delaneau, and Jonathan Marchini. Haplotype estimation for biobank-scale data sets. Nature genetics, 48(7):817, 2016.

[10] Trevor Hastie, Robert Tibshirani, and Jerome Friedman. The elements of statistical learning: data mining, inference, and prediction. Springer Science & Business Media, 2009.

[11] Ziad Al Bkhetan, Justin Zobel, Adam Kowalczyk, Karin Verspoor, and Benjamin Goudey. Exploring effective approaches for haplotype block phasing. BMC bioinformatics, 20(1):540, 2019.

[12] Yongwook Choi, Agnes P Chan, Ewen Kirkness, Amalio Telenti, and Nicholas J Schork. Comparison of phasing strategies for whole human genomes. PLoS genetics, 14(4):e1007308, 2018.

[13] Sharon R Browning and Brian L Browning. Haplotype phasing: existing methods and new developments. Nature Reviews Genetics, 12(10):703–714, 2011.

[14] Amy L Williams, Nick Patterson, Joseph Glessner, Hakon Hakonarson, and David Reich. Phasing of many thousands of genotyped samples. The American Journal of Human Genetics, 91(2):238–251, 2012.

[15] Omer Sagi and Lior Rokach. Ensemble learning: A survey. Wiley Interdisciplinary Reviews: Data Mining and Knowledge Discovery, 8(4):e1249, 2018.

[16] Olivier Delaneau, Jonathan Marchini, and Jean-François Zagury. A linear complexity phasing method for thousands of genomes. Nature methods, 9(2):179–181, 2012.

[17] International HapMap 3 Consortium et al. Integrating common and rare genetic variation in diverse human populations. Nature, 467(7311):52, 2010.

[18] Christopher C Chang, Carson C Chow, Laurent CAM Tellier, Shashaank Vattikuti, Shaun M Purcell, and James J Lee. Second-generation plink: rising to the challenge of larger and richer datasets. Gigascience, 4(1):7, 2015.

[19] Po-Ru Loh, Pier Francesco Palamara, and Alkes L Price. Fast and accurate long-range phasing in a uk biobank cohort. Nature genetics, 48(7):811–816, 2016.

[20] Anthony Francis Herzig, Teresa Nutile, Marie-Claude Babron, Marina Ciullo, Céline Bellenguez, and Anne-Louise Leutenegger. Strategies for phasing and imputation in a population isolate. Genetic epidemiology, 2018.

[21] Shefali S Verma, Mariza De Andrade, Gerard Tromp, Helena Kuivaniemi, Elizabeth Pugh, Bahram Namjou-Khales, Shubhabrata Mukherjee, Gail P Jarvik, Leah C Kottyan, Amber Burt, et al. Imputation and quality control steps for combining multiple genome-wide datasets. Frontiers in genetics, 5:370, 2014.

[22] Jonathan Marchini, David Cutler, Nick Patterson, Matthew Stephens, Eleazar Eskin, Eran Halperin, Shin Lin, Zhaohui S Qin, Heather M Munro, Gonçalo R Abecasis, et al. A comparison of phasing algorithms for trios and unrelated individuals. The American Journal of Human Genetics, 78(3):437–450, 2006.

[23] Cathy C Laurie, Kimberly F Doheny, Daniel B Mirel, Elizabeth W Pugh, Laura J Bierut, Tushar Bhangale, Frederick Boehm, Neil E Caporaso, Marilyn C Cornelis, Howard J Edenberg, et al. Quality control and quality assurance in genotypic data for genome-wide association studies. Genetic epidemiology, 34(6):591–602, 2010.

[24] Pierre Faux, Pierre Geurts, and Tom Druet. A random forests framework for modeling haplotypes as mosaics of reference haplotypes. Frontiers in genetics, 10:562, 2019.

