## Supplementary Materials for "Evaluation of consensus strategies for haplotype phasing"

Ziad Al Bkhetan, Gursharan Chana,
Kotagiri Ramamohanarao, Karin Verspoor, and Benjamin Goudey

**3 Supplementary tables** **5**

**4 Supplementary Figures** **9**

**5 Links for online resource** **15**

### 1 Supplementary Methods

#### 1.1 Dataset details

The data used in this study and not detailed in the main manuscript in described below:

1. **Reference dataset:** The reference panel for genotype imputation was formed purely from the HRC dataset by excluding 8,189 samples used in the Hap+HRC dataset and all samples from the 1000 genome project as they are from different populations.

2. **Low-density datasets**: Five different datasets were generated from chromosomes 16 and 21 of the Hap+HRC datasets. PLINK was applied to remove a random SNP from each pair if the distance between them is less than the desired resolution. The used densities were 1 SNP per 1kb, 2kb, 3kb, 4kb and 5kb.

3. **Different population size datasets**: The datasets with different population sizes were generated by a random selection of the individuals from the Hap+HRC dataset. Five sample sizes were generated as follows: 500, 1000, 2000, 4000, 6000 and all individuals 8,258 for chromosome 16 and 21.

4. **Different ethnicity dataset**: This dataset was generated purely from HapMap III datasets for chromosme 21 and the same SNPs as the chromosme 21 in the Hap+HRC dataset mentioned above. Parents genotype data were excluded but used to resolve children's haplotypes for evaluation purposes. The final dataset contains 838 individuals as follows: African ancestry in Southwest USA (ASW): 43, Utah residents with Northern and Western European ancestry from the CEPH collection (CEU): 69, Han Chinese in Beijing, China (CHB): 84, Chinese in Metropolitan Denver, Colorado (CHD): 85, Gujarati Indians in Houston, Texas (GIH): 88, Japanese in Tokyo, Japan (JPT): 86, Luhya in Webuye, Kenya (LWK): 90, Mexican ancestry in Los Angeles, California (MXL): 27, Maasai in Kinyawa, Kenya (MKK ): 115, Toscani in Italia (TSI): 88, and Yoruba in Ibadan, Nigeria (YRI): 63.

Details about the datasets used in this study are summarised in Table 1. Genetic maps of the reference genome hg19 were obtained from public resources and used with the tools for phasing and imputation when required. The HRC dataset is available through the European Genome-phenome Archive (dataset reference: EGAD00001002729). See Links below for data online resources.

Table 1 Details of datasets used in this study after preparation. Chromosomes' columns represent SNP count within each chromosome.

| Dataset | Samples | Chr2 | Chr6 | Chr11 | Chr16 | Chr21 |
|---|---|---|---|---|---|---|
| **Hap+HRC** | 8,258 | 50,521 | 38,930 | 30,580 | 21,928 | 9,319 |
| **Reference** | 16,481 | 3,392,237 | 2,460,111 | 1,936,990 | 1,281,297 | 531,276 |
| **HapMap-CEU** | 52 | 114,897 | 89,850 | 69,852 | 43,798 | 19,056 |

#### 1.2 Consensus estimator construction

Consensus estimators of all possible combinations of three tools out of SHAPEIT2, SHAPEIT3, EAGLE2, and HAPI-UR were constructed and evaluated. We have shown previously that adding more tools does not improve accuracy if the added tool is substantially less accurate than the ones currently being used [1]. Therefore, we limited the construction of our consensus estimator to only three different tools, shown to have the highest accuracies.

When constructing a consensus estimator from multiple iterations of non-deterministic tool, we evaluated different combinations of the results. With respect to Hap+HRC dataset, the estimator constructed from all possible combinations of 3, 5, 7, 9 out of 10 different iterations of the same tool. For the consensus of a large number of iterations (50), we limited the results to 15 random combinations for each odd number from 3 to 49 out of 50.

We use a consistent naming convention to describe a particular implementation of consHap throughout this work. Individual tools are abbreviated ($S_2$: SHAPEIT2, $S_3$: SHAPEIT3, $E_2$: EAGLE2, $H_r$: HAPI-UR) and concatenated together if multiple tools are used. If multiple outputs from a single tool are used, we denote the number of iterations in parentheses. Thus consHap-$S_2E_2S_3$ is based on the output of SHAPEIT2, EAGLE2, and SHAPEIT3 while consHap-$S_2$*(3)* is based on three different iterations of SHAPEIT2.

##### 1.3 Evaluation of haplotype phasing

The haplotypes of 52 children from CEU cohort were resolved using their parental genotypes. Transmission phasing was able to resolve ∼75% of child heterozygous SNPs when at least one parent has a homozygous SNP in the same locus. We clarify that the parents' genotype data were excluded from the dataset used in this study as the focus is on phasing unrelated individuals. The same procedure was applied to all children from different populations to evaluate phasing accuracy on different ethnicity.

Phasing evaluation was conducted through switch error (SE) calculated for the 52 samples mentioned above similarly to these studies [1, 2, 3]. The average switch errors of the individuals are averaged and reported for the whole dataset. Missing SNPs, SNPs with Mendel errors, and unresolved SNPs (Child's SNP where both parents have heterozygous SNP within the same locus) were excluded from the evaluation.

##### 1.4 Evaluation of genotype imputation

Hap+HRC datasets were aligned to the reference panel datasets (The same alternate and reference allele for each common SNP between the reference and Hap+HRC dataset) which is mandatory for both Beagle5 and pbwt. Minimac3 seems to account for that as its results were not affected by the alignment. Imputation tools were applied to all phased chromosomes obtained from phasing experiments (by individual tools and consensus estimators). Imputation tools were applied using their default parameters. Genetic maps were only provided to Beagle5 as they are required for this tool. Reference panel was converted into pbwt format when provided to pbwt tool.

Imputation evaluation was carried out on 8,189 individuals from both UK10UK and IBD cohorts with respect to all SNPs in HRC dataset that do not belong to the SNPs of the Hap+HRC study (similarly to masked analysis concept [4]). Squared Pearson correlation $r^2$ as defined in Eq 1 was calculated for the real genotype and both the sum of the posterior allele probabilities (${r_a}^2$), and the imputed genotype (${r_g}^2$) for each SNP separately, similarly to the evaluations carried out previously [5, 6, 7, 8]. We clarify that $r^2$ is considered zero when the output of the imputation tool does not change for a single SNP across all individuals, as the correlation can not be determined in this case.

$$r_{(s,s')} = cor(s, s') = \frac{\sum_{i=1}^{n} (s_i - \bar{s})({s'}_i - \bar{s'})}{\sqrt{\sum (s_i - \bar{s})^2 \sum ({s'}_i - \bar{s'})^2}} \tag{1}$$

Where: $s$ represents the value of the real genotype of a SNP as 0, 1, or 2. $s'$: represents the imputed genotype of a SNP in two possible forms: either a value of 0, 1, or 2 similar to the real genotypes representation when calculating ${r_g}^2$ or a value in the range (0, 2) as the sum of the posterior allele probabilities provided by the tools when calculating ${r_a}^2$. All evaluation metrics were averaged for all SNPs in three MAF ranges similar to the ranges in the study [6]: 0.01% to 0.5%, 0.5% to 5%, and 5% to 50%.

##### 1.5 Correlation assessment of phasing and imputation accuracy

This analysis was carried out on 52 individuals from the HapMap project where their true haplotypes (resolved using family information) and true genotypes for the imputed SNPs are available. Both imputation and phasing accuracies were calculated at an individual scale for the five chromosomes. Imputation accuracy was measured as the percentage of the correctly imputed SNPs as a proportion of all individual's imputed SNPs. Only the common SNPs in HapMap III and HRC datasets that do not belong to the SNPs in the Hap+HRC dataset were assessed. The counts of the common SNPs are reported in the third row in Table 3. Phasing accuracy was calculated as (100 - Switch error %) also at an individual scale. We used a Pearson correlation to assess the relationship between each pair of phasing and imputation tools, with reported significance at a p-value threshold = 0.05.

#### 2 Supplementary experiments

##### 2.1 Evaluation at individual scale

Switch error results are averaged for switch errors calculated for the individuals in the dataset. While the consensus estimators reduced the overall switch error, there were some individuals in the dataset phased more accurately by one tool, or one iteration of a non-deterministic tool compared to the consensus estimator. The switch error of consHap-$S_2E_2S_3$ and consHap-$S_2(3)$ for such cases was very similar to the most accurate tool or iteration (0.065% and 0.066% average difference, respectively) compared to the average difference between the most accurate two tools or iterations (0.2% and 0.1%, respectively). Table 2 shows the percentage of individuals that are phased by the tool in the related row compared to the tool in the related column. For example, 63.5 (row:1, column:1) means that 63.5% of all individuals were phased more accurately by the consensus consHap-$S_2E_2S_3$ compared to eh consensus consHap-$S_2(3)$. As we represent only the percentage of individuals that are phased more accurately by a tool compared to another one, the percentage of individuals that are phased similarly can be calculated from each two transposed cells. For example, the percentage of the individual phased with equal accuracy between consHap-$S_2E_2S_3$ and consHap-$S_2(3)$ is 100 - (Table2[consHap-$S_2E_2S_3$, consHap-$S_2(3)$] + Table2[consHap-$S_2(3)$, consHap-$S_2E_2S_3$]) = 100 - 63.5 - 31.2 = 0.3%.

Table 2 also highlights the performance of SHAPEIT2 and EAGLE2 individually, showing that 59% of CEU samples were phased more accurately by SHAPEIT2 compared to EAGLE2. HAPI-UR did not phase any individual more accurately than SHAPEIT2 or EAGLE2, but it did for 3% of the individuals compared to SHAPEIT3.

Table 2 Comparison of tool performance at an individual scale. Numbers represent the percentage of the individuals phased more accurately by the tool in the related row compared to the tool in the related column. Total number of individuals used to generate this table is 52 individuals for 5 chromosomes = 260 samples.

| – | **consHap-$S_2E_2S_3$** | **consHap-$S_2(3)$** | **SHAPEIT2** | **EAGLE2** | **SHAPEIT3** | **HAPI-UR** |
|---|---|---|---|---|---|---|
| **consHap-$S_2E_2S_3$** | 0.0 | 63.5 | 84.6 | 85.0 | 96.9 | 100.0 |
| **consHap-$S_2(3)$** | 31.2 | 0.0 | 75.4 | 73.1 | 93.5 | 100.0 |
| **SHAPEIT2** | 11.9 | 18.1 | 0.0 | 58.8 | 83.5 | 99.6 |
| **EAGLE2** | 12.3 | 23.1 | 39.2 | 0.0 | 75.8 | 99.6 |
| **SHAPEIT3** | 1.5 | 4.2 | 15.4 | 21.5 | 0.0 | 96.9 |
| **HAPI-UR** | 0.0 | 0.0 | 0.0 | 0.0 | 3.1 | 0.0 |

#### 2.2 Performance evaluation

The performance evaluation was assessed based on the running time and memory usage during the execution of all phasing and imputation tools. We used a server of 32 VCPU, 2300 MHz and 64GB RAM operated by Ubuntu 16.04 LTS (Xenial). Tools applied with default parameters or specifying explicitly one thread for execution if this option is available. We don't report evaluation for all scenarios due to the need to do phasing and imputation numerous times and in parallel using different thread count based on the available resources at a particular time.

Different manipulated versions of chromosome 21 dataset (the largest version was 19.1MB in plink format) were used for phasing evaluation. The same dataset phased by consHap-$S_2E_2S_3$ (23.6MB compressed VCF) in addition to chr21 reference panel (531,276 SNPs of 16,481 individuals, 534.5MB compressed VCF) were used to assess the performance of imputation tools.

The total time required to construct a consensus estimator is either the sum of the performance time for each tool when the tools are applied sequentially or the performance time of the slowest tool when the tools are applied in parallel in addition to the time required to aggregate phased haplotypes that can be negligible compared to the performance time of phasing tools. Constructing consHap-$S_2E_2S_3$ on chromosome 21 of the Hap+HRC dataset (large size and high density) required 32 hours. Phasing chromosome 21 of 500 individuals using SHAPEIT2 (the slowest tool in this study) required 40 minutes, therefore, the execution time of consHap-S2($n$) is 40 * $n$. However, these results represent the worst-case scenario where phasing is applied sequentially and using one thread.

The maximum RAM usage had a similar pattern to the performance time except for SHAPEIT3 that used more than SHAPEIT2. Phasing 9,319 SNPs of 8,258 individuals required less than 1 GB for HAPI-UR and EAGLE2, and 2 and 3 GB for SHAPEIT2 and SHAPEIT3 respectively.

Regarding genotype imputation performance, pbwt was the fastest tool accomplishing the imputation of the whole chromosome 21 within 40 minutes, 2 times faster than Beagle5 which required 77 minutes while Minimac3 spent 19 hours. pbwt was the optimal tool when considering memory usage. Beagle5 used $\sim$40GB of the ram during its execution. SHAPEIT2, SHAPEIT3 and Beagle5 can run using multiple threads, while Minimac3 requires another version. We have noticed after finalising the experiments that Minimac4 is available now, and it is claimed to be at least 2 times faster than Minimac3.

#### 3 Supplementary tables

Table 3 A comparison of correctly phased haplotype blocks. BL: the average length of the correctly phased haplotype blocks at an individual scale. BC the average count of different haplotype blocks at an individual scale (equivalent to switch count). This table represents the reflection of the accuracy improvement obtained by the consensus estimator on increasing the length of the correctly phased haplotype blocks. The average length of correctly phased haplotype blocks obtained by cons-S2E2S3 and cons-S2(3) is 421, and 403 SNPs respectively, representing improvements of 12% and 7.3% compared to the length of the correctly phased blocks obtained by the best individual tool in this test (∼40 SNPs longer than the most accurate individual tool). Best results are in bold. Best results obtained by individual tool are underlined. The length of the correctly phased haplotype block was calculated at an individual scale as the SNP count between two consecutive switch errors occurred in heterozygous SNPs.

| **Phasing** | **Chr2** | | **Chr6** | | **Chr11** | | **Chr16** | | **Chr21** | |
|---|---|---|---|---|---|---|---|---|---|---|
| | **BL** | **BC** | **BL** | **BC** | **BL** | **BC** | **BL** | **BC** | **BL** | **BC** |
| **consHap-**$S_2E_2S_3$ | **466** | **113** | **494** | **83** | **457** | **73** | **358** | **63** | **331** | **29** |
| **consHap-**$S_2(3)$ | 450 | 116 | 470 | 85 | 439 | 76 | 356 | 65 | 302 | 31 |
| **SHAPEIT2** | <u>418</u> | <u>125</u> | <u>437</u> | <u>92</u> | <u>413</u> | <u>80</u> | <u>329</u> | <u>69</u> | 284 | <u>33</u> |
| **EAGLE2** | 398 | 129 | 424 | 94 | 392 | 83 | 314 | 71 | <u>287</u> | 34 |
| **SHAPEIT3** | 361 | 144 | 383 | 107 | 354 | 93 | 284 | 79 | 272 | 36 |
| **HAPI-UR** | 258 | 198 | 279 | 143 | 249 | 128 | 207 | 108 | 187 | 51 |

Table 4 Switch error % obtained by consensus estimator constructed from a combination of three tools (out of 4) calculated across 5 chromosomes. $S_2$: SHAPEIT2, $S_3$: SHAPEIT3, $E_3$: EAGLE2, and $H_r$: HAPI-UR. Consensus estimators are sorted based on the average switch error (lowest is first). Numbers in bold are the lowest switch errors.

| **Consensus Approach** | **Chr2** | **Chr6** | **Chr11** | **Chr16** | **Chr21** |
|---|---|---|---|---|---|
| **consHap-**$S_2E_2S_3$ | **0.859** | **0.825** | **0.921** | **1.103** | **1.233** |
| **consHap-**$S_2E_2H_r$ | 0.875 | 0.836 | 0.947 | 1.114 | 1.270 |
| **consHap-**$S_2H_rS_3$ | 0.897 | 0.873 | 0.969 | 1.150 | 1.317 |
| **consHap-**$S_3E_2H_r$ | 0.914 | 0.874 | 0.971 | 1.171 | 1.309 |

Table 5 Switch error % obtained by consensus estimator constructed from a combination of multiple iterations of different tools (out of 4) calculated across 5 chromosomes. S2: SHAPEIT2, S3: SHAPEIT3, and E2: EAGLE2. Numbers between parenthesis represent the number of iterations of the associated tool. Consensus estimators are sorted based on the average switch error (lowest error is first).

| **Consensus Approach** | **Chr16** | **Chr21** | **Mean-SE** |
|---|---|---|---|
| consHap-$S_2(8)S_3(0)E_2(7)$ | 0.992 | 1.122 | 1.057 |
| consHap-$S_2(7)S_3(0)E_2(6)$ | 0.995 | 1.131 | 1.063 |
| consHap-$S_2(6)S_3(0)E_2(5)$ | 1.002 | 1.135 | 1.068 |
| consHap-$S_2(5)S_3(0)E_2(4)$ | 1.010 | 1.135 | 1.073 |
| consHap-$S_2(4)S_3(0)E_2(3)$ | 1.015 | 1.141 | 1.078 |
| consHap-$S_2(7)S_3(0)E_2(8)$ | 1.010 | 1.155 | 1.082 |
| consHap-$S_2(6)S_3(0)E_2(7)$ | 1.018 | 1.162 | 1.090 |
| consHap-$S_2(5)S_3(0)E_2(6)$ | 1.020 | 1.173 | 1.097 |
| consHap-$S_2(4)S_3(0)E_2(5)$ | 1.032 | 1.175 | 1.104 |
| consHap-$S_2(7)S_3(7)E_2(7)$ | 1.038 | 1.174 | 1.106 |
| consHap-$S_2(3)S_3(0)E_2(2)$ | 1.035 | 1.185 | 1.110 |
| consHap-$S_2(5)S_3(5)E_2(5)$ | 1.045 | 1.176 | 1.111 |
| consHap-$S_2(3)S_3(0)E_2(4)$ | 1.041 | 1.184 | 1.112 |
| consHap-$S_2(3)S_3(3)E_2(3)$ | 1.047 | 1.178 | 1.112 |
| consHap-$S_2(2)S_3(0)E_2(3)$ | 1.050 | 1.198 | 1.124 |
| consHap-$S_2(1)S_3(0)E_2(2)$ | 1.090 | 1.231 | 1.161 |
| consHap-$S_2(2)S_3(0)E_2(1)$ | 1.104 | 1.224 | 1.164 |
| consHap-$S_2E_2S_3$ | 1.103 | 1.233 | 1.168 |

Table 6 Accuracy improvement when adding more iterations to the consensus construction. SE: the improvement in switch error. Switches: the reduction of switches. Numbers were calculated for the mean of switches, and switch errors for all combinations of the same number of iterations.

| **Iterations** | **Chromosome 16** | | **Chromosome 21** | |
|---|---|---|---|---|
| | **SE** | **Switches** | **SE** | **Switches** |
| **1 to 3** | 6.1% | 224 | 6.53% | 113 |
| **3 to 5** | 2.15% | 74 | 2.35% | 37 |
| **5 to 7** | 1.04% | 35 | 1.14% | 18 |
| **7 to 9** | 0.32% | 10 | 0.82% | 13 |

Table 7 ${r_a}^2$ and ${r_g}^2$ summarised for 37,442,564,700 SNPs with MAF in (0.0001, 0.005]. Without phasing, there were 37,322,838,335, 37,377,063,365, and 37,309,197,496 correctly imputed SNPs by Beagle5, Minimac3 and bwt, respectively.

| **Imputed by** | **Phased by** | ${r_a}^2$ | ${r_g}^2$ | **Accuracy** | **Improvement** |
|---|---|---|---|---|---|
| Beagle5 | consHap-$S_2(8)E_2(7)$ | 0.314 | 0.419 | 99.929 | 191,419 |
| Beagle5 | consHap-$S_2E_2S_3$ | 0.309 | 0.414 | 99.929 | 69934 |
| Beagle5 | consHap-$S_2(3)$ | 0.308 | 0.413 | 99.929 | 62303 |
| Beagle5 | EAGLE2 | 0.307 | 0.411 | 99.929 | 45799 |
| Beagle5 | SHAPEIT2 | 0.306 | 0.411 | 99.929 | 336900 |
| Beagle5 | SHAPEIT3 | 0.299 | 0.404 | 99.928 | 242223 |
| Beagle5 | HAPI-UR | 0.296 | 0.400 | 99.927 | 92611450 |
| Beagle5 | unphased | 0.066 | 0.137 | 99.688 | - |
| Minimac3 | consHap-$S_2(8)E_2(7)$ | 0.315 | 0.419 | 99.930 | 172308 |
| Minimac3 | consHap-$S_2E_2S_3$ | 0.310 | 0.414 | 99.930 | 66852 |
| Minimac3 | consHap-$S_2(3)$ | 0.309 | 0.413 | 99.929 | 39526 |
| Minimac3 | EAGLE2 | 0.308 | 0.412 | 99.929 | 52360 |
| Minimac3 | SHAPEIT2 | 0.307 | 0.411 | 99.929 | 304959 |
| Minimac3 | SHAPEIT3 | 0.299 | 0.404 | 99.928 | 194923 |
| Minimac3 | HAPI-UR | 0.297 | 0.400 | 99.928 | 38729491 |
| Minimac3 | unphased | 0.065 | 0.110 | 99.826 | - |
| pbwt | consHap-$S_2(8)E_2(7)$ | 0.279 | 0.382 | 99.922 | 242443 |
| pbwt | consHap-$S_2E_2S_3$ | 0.274 | 0.378 | 99.922 | 88760 |
| pbwt | consHap-$S_2(3)$ | 0.272 | 0.376 | 99.922 | 67560 |
| pbwt | EAGLE2 | 0.271 | 0.374 | 99.922 | 77275 |
| pbwt | SHAPEIT2 | 0.270 | 0.374 | 99.922 | 433473 |
| pbwt | SHAPEIT3 | 0.261 | 0.365 | 99.920 | 359676 |
| pbwt | HAPI-UR | 0.259 | 0.361 | 99.919 | 103500627 |
| pbwt | unphased | 0.047 | 0.100 | 99.654 | - |

Table 8 ${r_a}^2$ and ${r_g}^2$ summarised for 7,052,080,185 SNPs with MAF in (0.005,0.05]. Without phasing, there were 6,846,636,842 , 6,866,563,139, and 6,811,686,105 correctly imputed SNPs by Beagle5, Minimac3 and bwt, respectively.

| **Imputed by** | **Phased by** | ${r_a}^2$ | ${r_g}^2$ | **Accuracy** | **Improvement** |
|---|---|---|---|---|---|
| Beagle5 | consHap-$S_2(8)E_2(7)$ | 0.768 | 0.8589 | 99.395 | 502812 |
| Beagle5 | consHap-$S_2E_2S_3$ | 0.766 | 0.857 | 99.387 | 160331 |
| Beagle5 | consHap-$S_2(3)$ | 0.765 | 0.856 | 99.385 | 356462 |
| Beagle5 | SHAPEIT2 | 0.763 | 0.855 | 99.379 | 27218 |
| Beagle5 | EAGLE2 | 0.763 | 0.855 | 99.380 | 992514 |
| Beagle5 | SHAPEIT3 | 0.758 | 0.852 | 99.364 | 1635093 |
| Beagle5 | HAPI-UR | 0.751 | 0.847 | 99.341 | 160479842 |
| Beagle5 | unphased | 0.327 | 0.512 | 97.087 | - |
| Minimac3 | consHap-$S_2(8)E_2(7)$ | 0.764 | 0.855 | 99.386 | 504664 |
| Minimac3 | consHap-$S_2E_2S_3$ | 0.762 | 0.854 | 99.379 | 134018 |
| Minimac3 | consHap-$S_2(3)$ | 0.761 | 0.853 | 99.377 | 341527 |
| Minimac3 | SHAPEIT2 | 0.759 | 0.852 | 99.372 | 51177 |
| Minimac3 | EAGLE2 | 0.759 | 0.852 | 99.372 | 1000277 |
| Minimac3 | SHAPEIT3 | 0.754 | 0.849 | 99.356 | 1663102 |
| Minimac3 | HAPI-UR | 0.746 | 0.843 | 99.332 | 140092807 |
| Minimac3 | unphased | 0.298 | 0.454 | 97.369 | - |
| pbwt | consHap-$S_2(8)E_2(7)$ | 0.739 | 0.839 | 99.305 | 576712 |
| pbwt | consHap-$S_2E_2S_3$ | 0.736 | 0.837 | 99.296 | 207810 |
| pbwt | consHap-$S_2(3)$ | 0.735 | 0.836 | 99.293 | 368865 |
| pbwt | EAGLE2 | 0.734 | 0.835 | 99.289 | 43738 |
| pbwt | SHAPEIT2 | 0.733 | 0.835 | 99.287 | 1173304 |
| pbwt | SHAPEIT3 | 0.727 | 0.831 | 99.270 | 1983544 |
| pbwt | HAPI-UR | 0.719 | 0.824 | 99.241 | 188778197 |
| pbwt | unphased | 0.249 | 0.427 | 96.601 | - |

Table 9 ${r_a}^2$ and ${r_g}^2$ summarised for 9,735,689,186 SNPs with MAF in (0.05,0.5]. Without phasing, there were 8,881,462,946 , 8,857,189,955, and 8,805,437,448 correctly imputed SNPs by Beagle5, Minimac3 and bwt, respectively.

| **Imputed by** | **Phased by** | ${r_a}^2$ | ${r_g}^2$ | **Accuracy** | **Improvement** |
|---|---|---|---|---|---|
| Beagle5 | consHap-$S_2(8)E_2(7)$ | 0.971 | 0.983 | 99.218 | 256652 |
| Beagle5 | consHap-$S_2E_2S_3$ | 0.971 | 0.984 | 99.210 | 330655 |
| Beagle5 | consHap-$S_2(3)$ | 0.971 | 0.984 | 99.206 | 277583 |
| Beagle5 | EAGLE2 | 0.971 | 0.984 | 99.204 | 235839 |
| Beagle5 | SHAPEIT2 | 0.971 | 0.983 | 99.200 | 1460372 |
| Beagle5 | SHAPEIT3 | 0.970 | 0.983 | 99.185 | 1862852 |
| Beagle5 | HAPI-UR | 0.969 | 0.983 | 99.163 | 779068641 |
| Beagle5 | unphased | 0.745 | 0.853 | 91.177 | - |
| Minimac3 | consHap-$S_2(8)E_2(7)$ | 0.970 | 0.983 | 99.197 | 796271 |
| Minimac3 | consHap-$S_2E_2S_3$ | 0.970 | 0.983 | 99.188 | 344538 |
| Minimac3 | consHap-$S_2(3)$ | 0.970 | 0.983 | 99.184 | 227954 |
| Minimac3 | EAGLE2 | 0.970 | 0.983 | 99.183 | 284205 |
| Minimac3 | SHAPEIT2 | 0.970 | 0.983 | 99.178 | 1531159 |
| Minimac3 | SHAPEIT3 | 0.969 | 0.983 | 99.162 | 1698946 |
| Minimac3 | HAPI-UR | 0.969 | 0.982 | 99.143 | 801593644 |
| Minimac3 | unphased | 0.759 | 0.855 | 90.943 | - |
| pbwt | consHap-$S_2(8)E_2(7)$ | 0.968 | 0.982 | 99.139 | 863115 |
| pbwt | consHap-$S_2E_2S_3$ | 0.968 | 0.982 | 99.129 | 426841 |
| pbwt | consHap-$S_2(3)$ | 0.968 | 0.982 | 99.124 | 149534 |
| pbwt | EAGLE2 | 0.968 | 0.982 | 99.124 | 420757 |
| pbwt | SHAPEIT2 | 0.968 | 0.982 | 99.118 | 1674540 |
| pbwt | SHAPEIT3 | 0.967 | 0.981 | 99.100 | 1814525 |
| pbwt | HAPI-UR | 0.966 | 0.981 | 99.079 | 848101462 |
| pbwt | unphased | 0.725 | 0.837 | 90.425 | - |

Table 10 Correlation assessment for all combination of phasing and imputation tools averaged for chromosomes 21, 16, 11, 6, and 2. When evaluating genotype imputation accuracy on 8,189 HRC individuals (results are above and in the manuscript), we use the switch error calculated for only the 52 HapMap III individuals (with known real haplotypes) as an indication of the quality of pre-phasing for the whole dataset (HapMap III and HRC individuals combined in the Hap+HRC dataset). For more accurate evaluation of the impact of phasing on the genotype imputation, we assess the correlation between phasing accuracy and genotype accuracy of the same individual. The lack of known haplotype data limited this experiments to 52 individuals from HapMap III dataset (Phasing evaluation can be conducted on them by comparing phased haplotypes to the one resolved using paternal genotypes).

| **Imputation tool** | **Phasing tool** | **Correlation** | **p-value** |
|---|---|---|---|
| Beagle5 | EAGLE2 | 0.422 | 1.93e-12 |
| Beagle5 | HAPI-UR | 0.486 | 1.58e-16 |
| Beagle5 | consHap-$S_2E_2S_3$ | 0.406 | 1.56e-11 |
| Beagle5 | SHAPEIT2 | 0.38 | 3.42e-10 |
| Beagle5 | consHap-$S_2(3)$ | 0.354 | 6.33e-9 |
| Beagle5 | SHAPEIT3 | 0.366 | 1.65e-9 |
| Minimac3 | EAGLE2 | 0.37 | 1.10e-9 |
| Minimac3 | HAPI-UR | 0.463 | 5.78e-15 |
| Minimac3 | consHap-$S_2E_2S_3$ | 0.34 | 2.48e-8 |
| Minimac3 | SHAPEIT2 | 0.317 | 2.43e-7 |
| Minimac3 | consHap-$S_2(3)$ | 0.285 | 3.64e-6 |
| Minimac3 | SHAPEIT3 | 0.316 | 2.52e-7 |
| pbwt | EAGLE2 | 0.31 | 4.29e-7 |
| pbwt | HAPI-UR | 0.399 | 3.75e-11 |
| pbwt | consHap-$S_2E_2S_3$ | 0.309 | 4.93e-7 |
| pbwt | SHAPEIT2 | 0.290 | 2.42e-6 |
| pbwt | consHap-$S_2(3)$ | 0.266 | 1.65e-5 |
| pbwt | SHAPEIT3 | 0.262 | 2.28e-5 |

#### 4 Supplementary Figures

Figure 1 The relation between switch error% and the the number of tool iterations used to construct a consensus estimator from a single tool. X-axis is number of iterations. Results are summarised for all possible combinations of 3, 5, 7 and 9 combinations (out of 10 iterations) of each tool. We highlight that, all these tools are unstable by default but EAGLE2. Therefore, we changed the order of individuals within the dataset randomly with each iteration of EAGLE2.

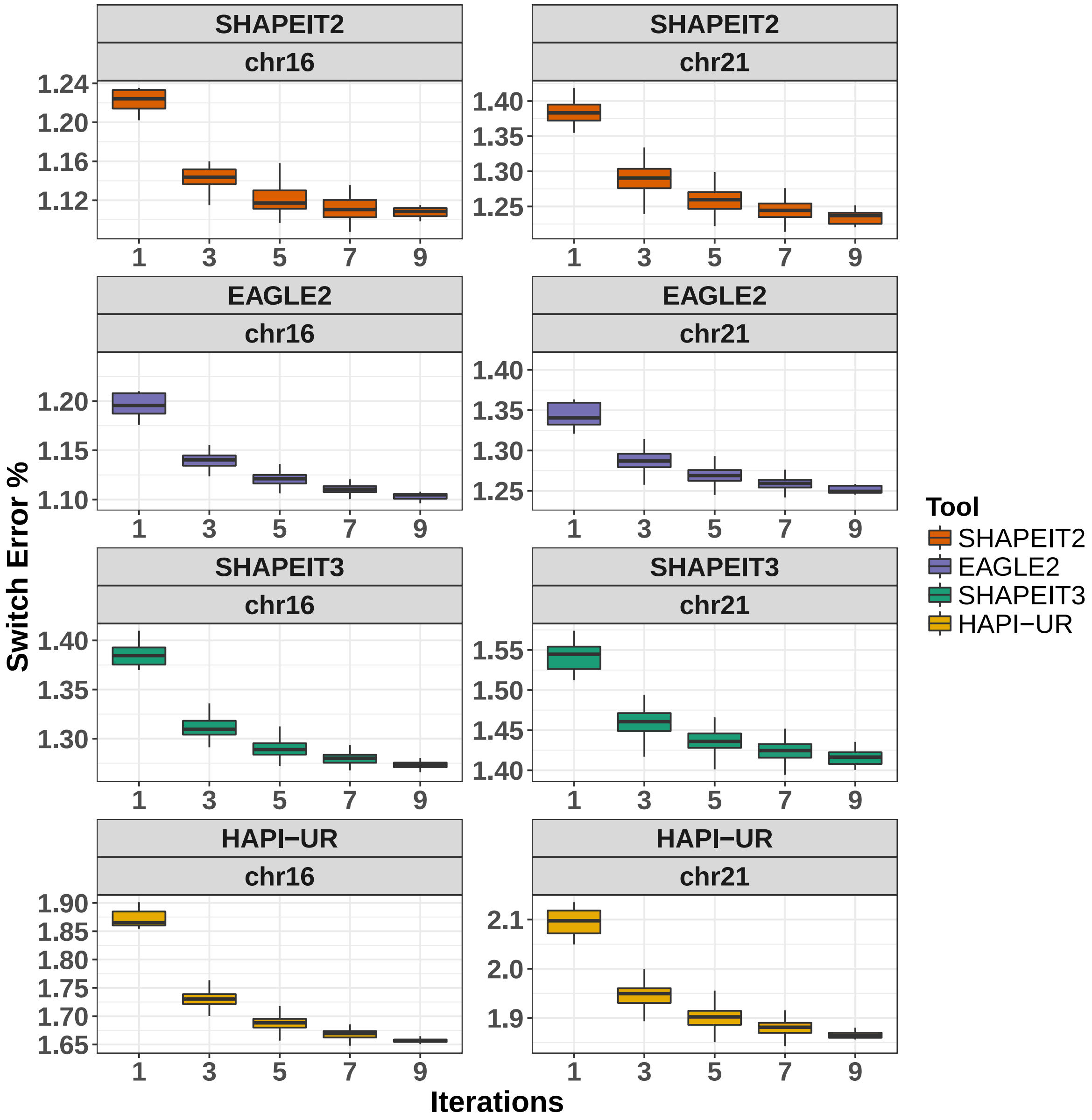

Figure 2 Comparison of the consensus approaches when applied to low and high quality dataset from chromosome 16. a) Comparison of the performance of the consHap-$S_2E_2S_3$ (pink horizontal line) and a consensus approach of several iterations of SHAPEIT2 (in brown) (1, 3, 5, 7 and 9) applied to large and high SNP density dataset (chromosome 16 as described in Hap+HRC dataset in the manuscript). b) Comparison of the performance of the consHap-$S_2E_2S_3$ and a consensus approach of several iterations of SHAPEIT2 for all odd numbers between 1 and 50. The data used in b) is a low quality version of chromosome 16 Hap+HRC study dataset, generated by reducing sample size to 500 individuals and SNP density by 50%.

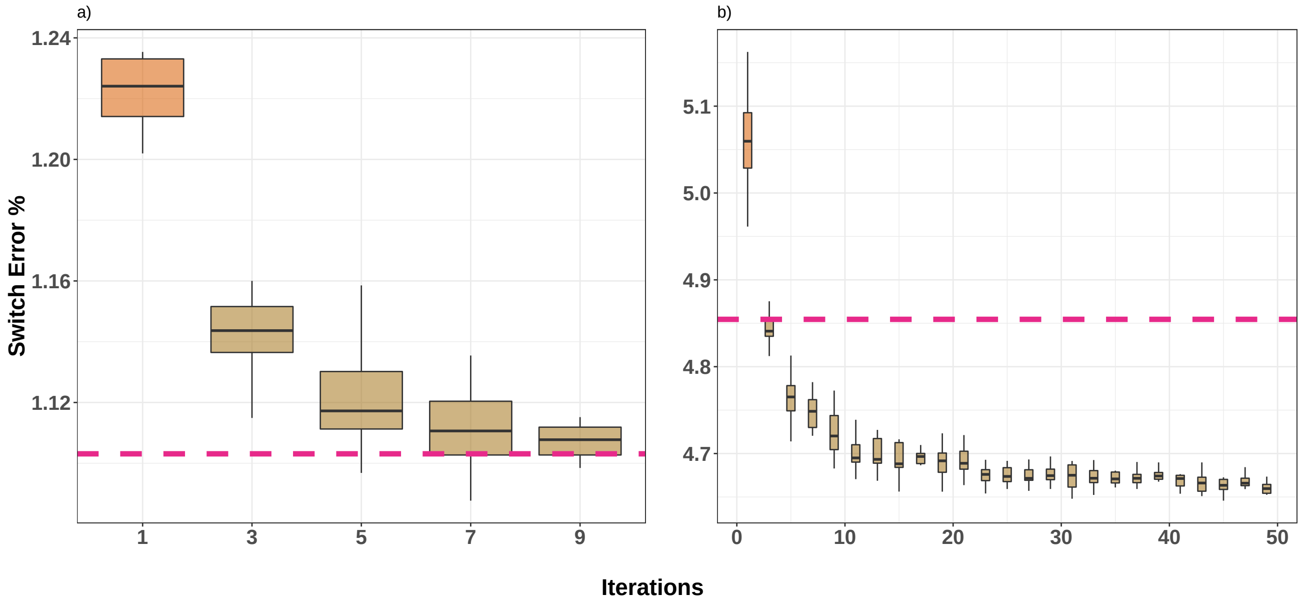

Figure 3 Switch error calculated for the consensus of SHAPEIT2, EAGLE2 and SHAPEIT3 (consHap-$S_2E_2S_3$) and the consensus of multiple iterations of SHAPEIT2 with respect to different population size and SNP density.

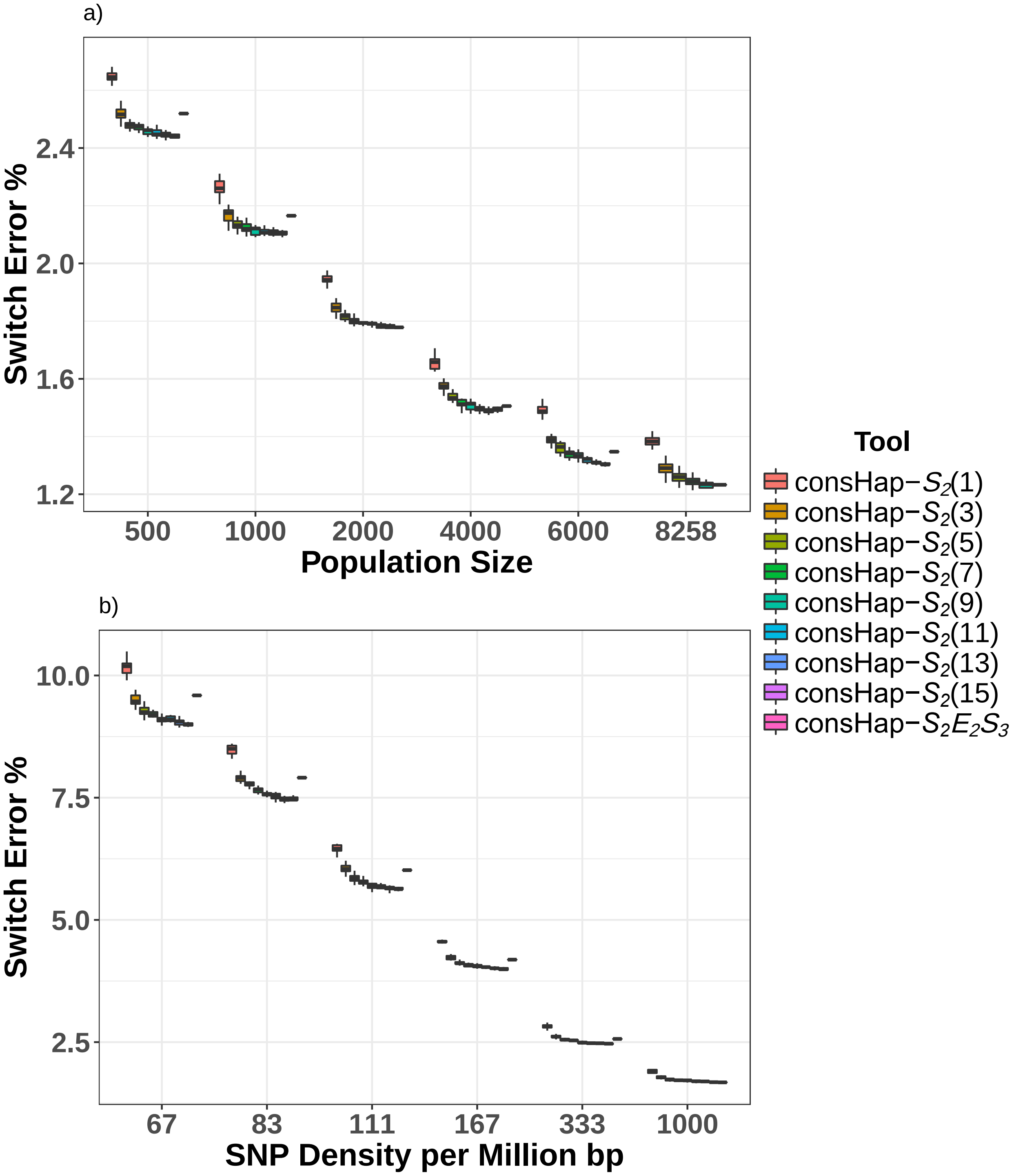

Figure 4 Comparison of the switch error obtained by consensus estimator approaches and individual tools when applied to low quality dataset. We clarify that both datasets contains only 500 individuals. SNP density in chromosome 16 was reduced to be 10,000 SNPs per the whole chromosomes. This figure shows that when EAGLE2 obtained high error rate, the consensus consHap-$S_2E_2S_3$ performs similarly to three iterations of SHAPEIT2. Adding more iterations of SHAPEIT2 improves the accuracy more than what can be obtained by three different tools.

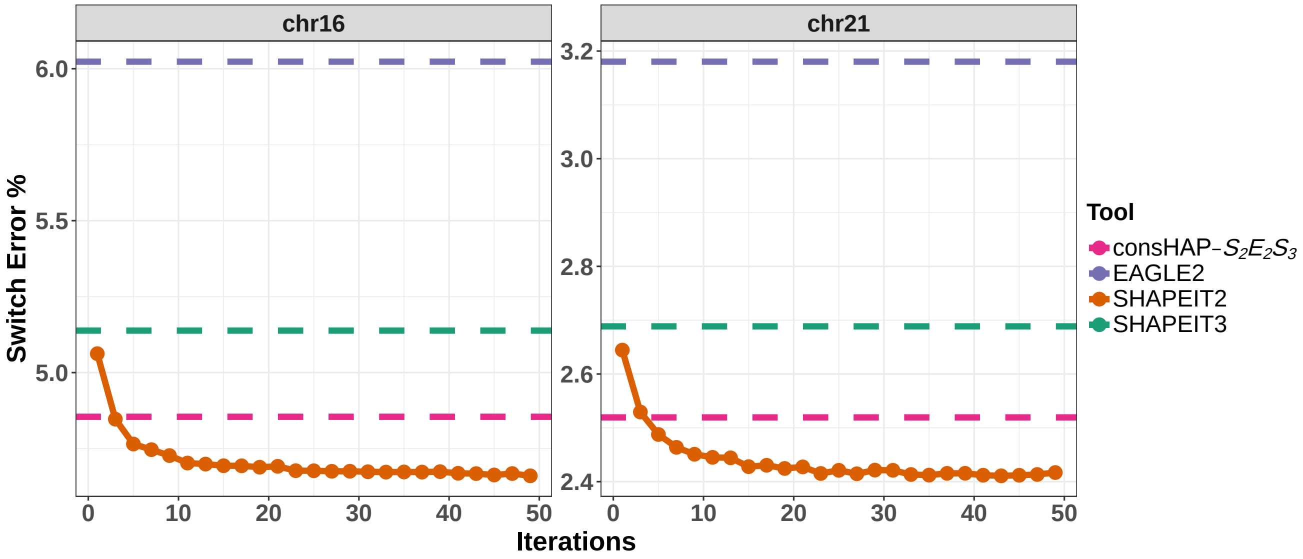

Figure 5 Comparison of the switch error obtained by a consensus estimator constructed by SHAPEIT2 and HAPI-UR for 1 to 49 iterations (only odd numbers). For each odd iteration ($i$), a 15 random combinations ($i$ iterations of 50 ) were used to construct a consensus of $i$ iterations. This figure shows that the accuracy of any consensus is highly associated with the tool used in its construction. 50 iterations of HAPI-UR doesn't outperform a single iteration of SHAPEIT2. Datasets used in this figure are described in the caption of Figure 4.

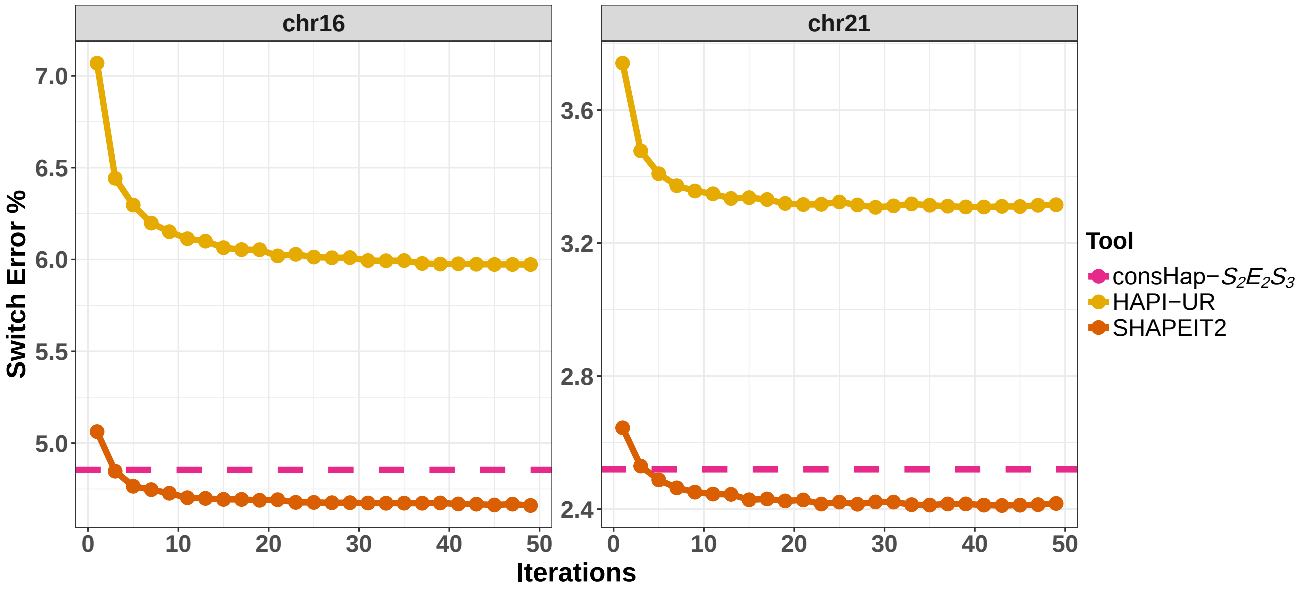

Figure 6 Correlation of phasing and imputation accuracy for 52 individuals from HapMap dataset. The x-axis represents phasing accuracy as 100 - Switch error %, while Y-axis represents the percentage of the correctly imputed SNP. Each point in the plots represents a pair of phasing and imputation accuracy for the same individual. The figure is generated from the results obtained from chromosomes 2, 6, 11, 16 and 21. The smoothed blue line represents a linear regression model of the imputation accuracy as a function of phasing accuracy. Each subplot is a combination of imputation and phasing tools. This figure is for the following phasing tools: consHap-$S_2E_2S_3$, SHAPEIT2, and EAGLE2.

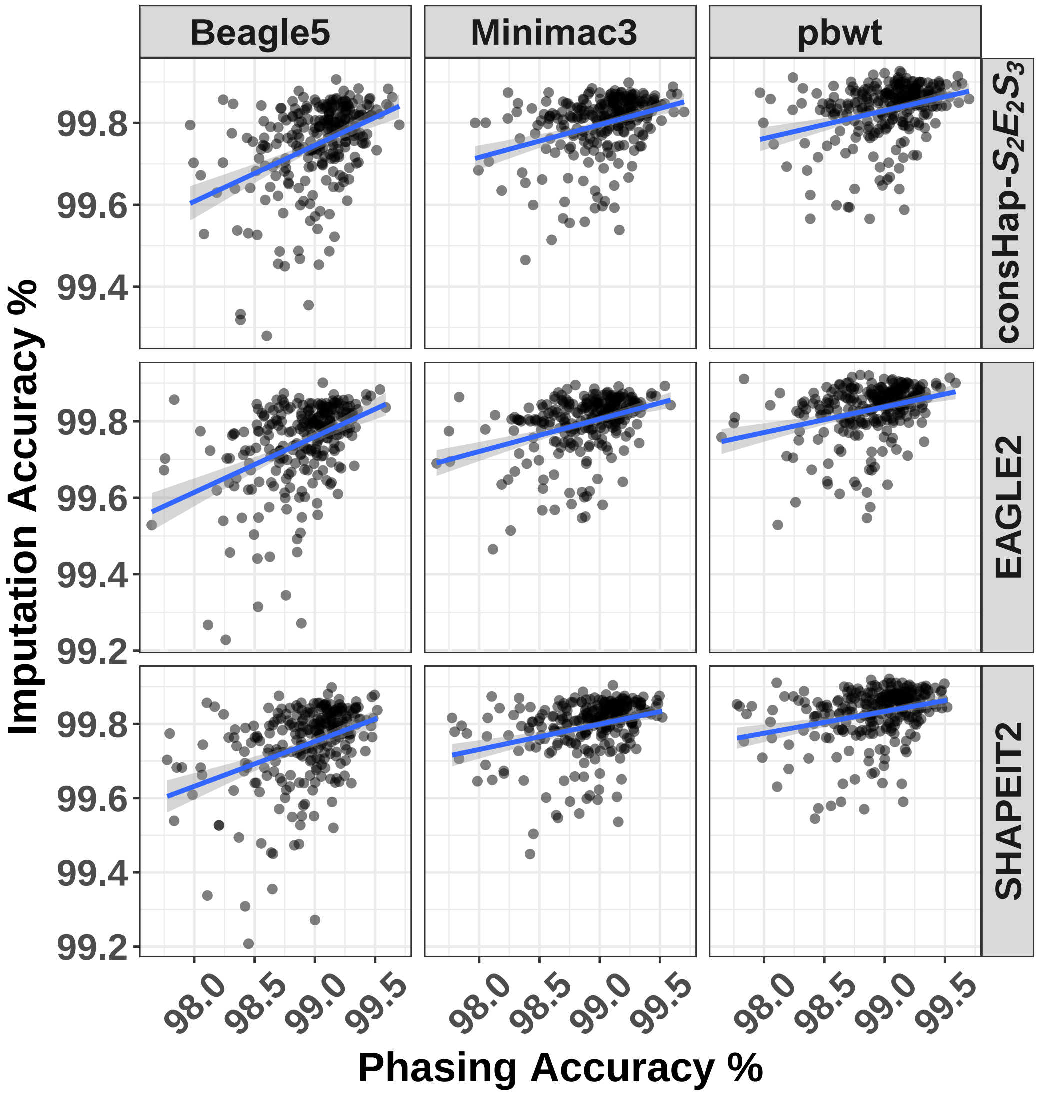

Figure 7 The same as Figure 6 but for the remaining phasing approaches: consHap-$S_2(3)$, SHAPEIT3 and HAPI-UR.

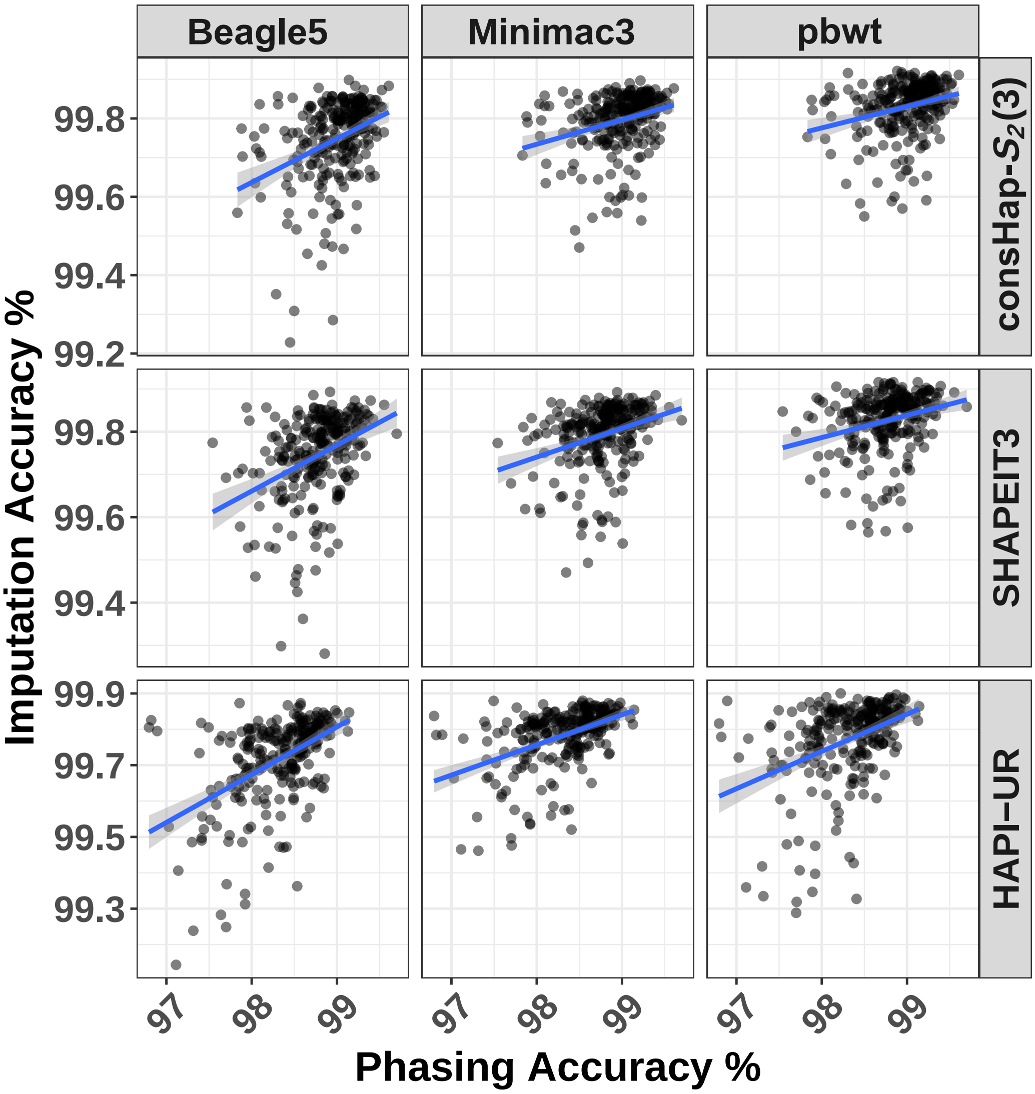

#### 5 Links for online resource

1. **consHap tool**: https://github.com/ziadbkh/consHap.
2. **SHAPEIT2**: https://mathgen.stats.ox.ac.uk/genetics_software/shapeit/shapeit.html.
3. **SHAPEIT3**: https://jmarchini.org/shapeit3/.
4. **EAGLE2**: https://data.broadinstitute.org/alkesgroup/Eagle.
5. **HAPI-UR**: https://code.google.com/archive/p/hapi-ur.
6. **Beagle5**: https://faculty.washington.edu/browning/beagle/beagle.html.
7. **Minimac3**: https://genome.sph.umich.edu/wiki/Minimac3.
8. **PBWT**: https://github.com/richarddurbin/pbwt.
9. **Genetic maps 1**: https://data.broadinstitute.org/alkesgroup/Eagle/downloads.
10. **Genetic maps 2**: http://bochet.gcc.biostat.washington.edu/beagle/genetic_maps.
11. **Haplotype Reference Consortium (HRC):** http://www.haplotype-reference-consortium.org.
12. **The HRC dataset:** Available through European Genome-phenome Archive (dataset reference: EGAD00001002729) at the link: https://www.ebi.ac.uk/ega/datasets/EGAD00001002729.
13. **HapMap III**: ftp://ftp.ncbi.nlm.nih.gov/HapMap/genotypes/HapMap3_r3.
14. **PLINK**: https://www.cog-genomics.org/plink2.
15. **Sanger imputation service**: https://imputation.sanger.ac.uk/.
16. **Michigan Imputation Server**: https://imputationserver.sph.umich.edu/index.html.

### References

[1] Z. Al Bkhetan, J. Zobel, A. Kowalczyk, K. Verspoor, and B. Goudey, “Exploring effective approaches for haplotype block phasing,” *BMC bioinformatics*, vol. 20, no. 1, p. 540, 2019.

[2] J. Marchini, D. Cutler, N. Patterson, M. Stephens, E. Eskin, E. Halperin, S. Lin, Z. S. Qin, H. M. Munro, G. R. Abecasis *et al.*, “A comparison of phasing algorithms for trios and unrelated individuals,” *The American Journal of Human Genetics*, vol. 78, no. 3, pp. 437–450, 2006.

[3] S. R. Browning and B. L. Browning, “Haplotype phasing: existing methods and new developments,” *Nature Reviews Genetics*, vol. 12, no. 10, pp. 703–714, 2011.

[4] S. S. Verma, M. De Andrade, G. Tromp, H. Kuivaniemi, E. Pugh, B. Namjou-Khales, S. Mukherjee, G. P. Jarvik, L. C. Kottyan, A. Burt *et al.*, “Imputation and quality control steps for combining multiple genome-wide datasets,” *Frontiers in genetics*, vol. 5, p. 370, 2014.

[5] B. L. Browning, Y. Zhou, and S. R. Browning, “A one-penny imputed genome from next-generation reference panels,” *The American Journal of Human Genetics*, vol. 103, no. 3, pp. 338–348, 2018.

[6] S. Das, L. Forer, S. Schönherr, C. Sidore, A. E. Locke, A. Kwong, S. I. Vrieze, E. Y. Chew, S. Levy, M. McGue *et al.*, “Next-generation genotype imputation service and methods,” *Nature genetics*, vol. 48, no. 10, p. 1284, 2016.

[7] B. Howie, C. Fuchsberger, M. Stephens, J. Marchini, and G. R. Abecasis, “Fast and accurate genotype imputation in genome-wide association studies through pre-phasing,” *Nature genetics*, vol. 44, no. 8, p. 955, 2012.

[8] Y. Li, C. Willer, S. Sanna, and G. Abecasis, “Genotype imputation,” *Annual review of genomics and human genetics*, vol. 10, pp. 387–406, 2009.
